# Nanoscale identification of organic pigments in biogenic calcite by atom probe tomography

**DOI:** 10.64898/2026.10.09.757918

**Authors:** Alexander Triccas, Tim M. Schwarz, Ya-Lan Hsu, Se-Ho-Kim, Christoph Freysoldt, Maïtena Dumont, Vaskar Sardhalia, Claudio Ferreira, Camille Chareyron, Marie Albéric, Baptiste Gault

**Affiliations:** Max Planck Institute for Sustainable Materials (formerly Max-Planck Institut für Eisenforschung GmbH), 40237 Düsseldorf, Germany; Department of Materials Science and Engineering, Korea University, Seoul 02841, Republic of Korea; Univ Rouen Normandie, CNRS, INSA Rouen Normandie, Groupe de Physique des Matériaux, UMR 6634, F-76000 Rouen, France; Sorbonne Université, CNRS, Laboratoire Chimie de la Matière Condensée de Paris, LCMCP, F-75005 Paris, France

**Keywords:** Atom probe tomography, Biomineralisation, Pigments, Nanostructure

## Abstract

Spatially-resolved mass spectrometry could contribute to the understanding of biological processes. Atom probe tomography (APT) is a technique which could, in principle, fill an analytical gap for nanoscale mass spectrometry. Yet, because of a limited understanding of the process of fragmentation of biomolecules, biological or biochemical interpretation of the obtained data has so far not been successful. Here, we report on analyses of synthetic and biogenic pigmented calcite minerals. The purple coloration of *Sphaerechinus granularis* sea urchin spines is due to the occlusion of polyhydroxylated naphthoquinone (PHNQ) molecules, mainly spinochrome A, within the calcite phase. Peaks relating to these pigments as well as to other intra-crystalline macromolecules were identified by APT analysis of the purple spine region with comparison to measurements of synthetic calcite single crystals, calcite crystals doped with PHNQ, and the uncoloured (white) region of sea urchin spines. To facilitate the molecular identification of peaks appearing solely in purple-coloured spines, which are possible fragments from PHNQ, we introduce novel APT data filtering techniques based on spatial and temporal correlations in the detection of ions of targeted mass-to-charge state ratios. This allows us to discuss the fragmentation of organic molecules under field conditions and improve the identification of peaks in the mass spectrum, while also determining nanoscale spatial distribution of fragments related to specifically targeted biomolecules in 3D and elucidate the relationship these organic pigment molecules have to the calcite lattice and inorganic elements such as magnesium. The study aims to improve identification of complex molecular structures by APT, including organic molecules, with the potential ability to provide structural information at a near-atomic scale for a range of materials.

## 1. Introduction

Biominerals are complex composites of inorganic and organic components organized across many scales and which perform functional roles to organisms across all seven kingdoms^1,2^. Some organic molecules, including proteins, polysaccharides and pigments from calcium carbonate (CaCO_3_) biominerals were shown to control *in vitro* several aspects of crystallisation, including nucleation and growth kinetics as well as crystal morphologies and polymorphism^3–13^. The extraction of these intracrystalline organic molecules also enables their thorough characterisation through e.g. amino acid composition^5,14^, SDS-PAGE electrophoresis^15^, FT-IR, mass and NMR spectroscopy^16^ as well as proteomics^17,18^. Their localisation and identification from the submicron down to the nano scale within biominerals, has been performed so far by a wide range of techniques, such as x-ray scanning scattering, diffraction, absorption spectroscopy and transmission spectromicroscopy as well as confocal Raman mapping, nanoscale secondary ion mass spectrometry, atomic force microscopy coupled with infrared or scanning transmission electron microscopy coupled with electron energy loss spectroscopy or energy-dispersive X-ray spectroscopy^16^. These powerful techniques are however mainly providing 2D information and 3D compositional information at the nanoscale is still missing. Therefore, there are still numerous questions pertaining to their intimate structure and composition within biominerals, which could provide significant insight into the biological mechanisms which control the formation of complex composite structures.

Atom probe tomography (APT) can perform 3D spatially-resolved mass spectrometry on a scale down to a nanometre, which is one to two orders of magnitude finer than other compositional techniques can typically achieve for biominerals^19,20^. Additional, APT can provide three-dimensional reconstruction of the analysed volume^21–23^. While APT has traditionally been applied to metals and semiconductors^24,25^, recent pioneering APT analyses of teeth in a range of species^26–28^, bones^29–32^ and CaCO_3_ shells^33–35^ have reported on the distribution of organic-rich regions within mineralised tissue, in addition to minor elements including Na, Mg, Fe and F. APT has also been conducted on several organic materials^36–42^ and biomolecules; either in dry form^43,44^, in cryogenically preserved aqueous solutions^45,46^, or as part of more complex biological entities including proteins^47,48^, viruses^49^ and sections of mammalian cells^50^.

In CaCO_3_ biominerals, peaks in the mass spectrum relating to organic molecules are identified as simple molecular fragments originating from the larger organic structure^51^. Assignment of each fragment to a specific chemical composition is complicated by organic molecules generating a large number of peaks in the mass spectrum which are usually identified as combinations of C, O, N and H^44^. Distinguishing between two fragment organic molecules at very close mass-to-charge state ratios can therefore be challenging as the mass resolving power of APT is limited below certain ranges. For example, predictable organic fragments CO_2_^+^ and CONH_2_^+^ separated by 0.023 Dalton (Da) will be detected at the same mass peak. Furthermore, the attribution of certain organic fragment peaks to a specific chemical group within the original molecular structure, such as the presence of acid/ester, alkene or amide groups for instance, cannot usually be achieved. The true nature of the bonding environments of peaks detected by APT remains unknown, as there can be rearrangement of the electrons and atoms within the fragment during the flight that can facilitate fragmentation of molecular ions and also the recombination of ions to generate new species under intense electric field conditions^52,53^. With only the mass-to-charge available, it is therefore difficult to correctly ascertain the chemical identity of each peak in the mass spectrum, and particularly relate this to individual organic molecules that are present in the larger structure.

Although fragmentation mechanisms in conventional mass spectrometry have been established through measurements of reference samples and advances in theory^54^, APT still lacks a framework to describe how organic molecules fragment in an intense electrostatic field. Comparison to peaks obtained in pure organic solution, either deposited on a field emitter or cryogenically frozen, may also present problems, as the intense electrostatic field can modify molecular orbitals and charge distributions^55^, alter bond strength and dissociation, and potentially change molecular conformation^52^, which will likely result in different fragmentation pathways and peaks detected in the mass spectrum. Since the electrostatic field is expected to modify the fragmentation process, the species ultimately detected are also likely to depend on specific analysis conditions (voltage vs. laser pulsing)^56^.

Wang and Kreuzer^55^ have suggested that fragmentation patterns of thiols under intense electric fields could be predicted by density-functional theory (DFT). More advanced quantum chemical calculations by Dietrich et al.^37^ were performed to simulate the fragmentation of n-tetradecane in an intense electrostatic field, yet their models do not fully agree with experimental data. These calculations predict ethyl and propyl cations, the smallest and most stable fragments in an organic compound, to be the most abundant species detected, whereas experimentally larger fragments are more dominant^36^. This discrepancy could arise from the bonding between a molecule and its surrounding environment, as simulations are performed on an isolated molecule free to deform in three-dimensions prior to or after fragmentation, which is not necessarily the case when the molecule is occluded within a crystalline material or cryogenically preserved.

Biominerals therefore provide an interesting case study as organic molecules are a constituent part of the overall composite material, where some are incorporated into the crystalline structure^57,58^. Of the calcite biominerals, the organic composition is well-studied in the spines of adult sea urchins^11,18,59^, which are Mg^60^ calcite-based biominerals. *Paracentrotus lividus* spines are reported to contain about 1.5 wt% of intracrystalline organics^61^ including proteins^5,8,18,59^, polysaccharides^59^, lipids and pigments^11,62^. The wide range of colours displayed by sea urchin spines has been attributed to the occlusion of polyhydroxylated naphthoquinone (PHNQ) molecules of about 250 Da^62–64^. While five main PHNQs have been reported for *P. lividus* (echinochrome A, spinochromes A, B, C and E)^63^, the purple coloration of their spines is specifically caused by the spinochrome A molecule (264 Da)^11,62^. The very high probability to find these low molecular weight molecules with a known structure within the coloured calcite sea urchin spines is a good test case to demonstrate the capability of APT to identify and track biomolecules, given that these molecules should be easier to interpret than complex proteins, which produce a large number of detected peaks in an APT mass spectrum^32^.

Identification of specific organic structures, like PHNQs, by APT can still be challenging as molecular ions are rarely observed, and fragmentation signatures generated under electrostatic field conditions are poorly understood. When a molecule undergoes field evaporation, all fragments are detected during a single pulse, forming what is referred to as a multiple event^65^. A mass spectrum of ions detected as part of multiple events exhibits lower background levels and can reveal additional mass peaks^66^. Ions on multiple events are typically detected in proximity of each other, usually within 3–4 mm on a straight flight path instrument^67,68^. Dissociation can lead to a kinetic-energy release (KER)^67,69,70^ and the intense Coulomb forces from the charged fragments in the vicinity of each other lead to a deflection of the two ions from their expected trajectories that can be measured by the distance between the two detector impact positions^67^. In addition, fragmentation can occur while the molecule is still attached to the surface^55^, which means that fragments of a single molecule may not be emitted by an individual pulse but rather in events that are detected in close succession. To ascertain this, filtering for these “proximal events” is also possible, e.g. ions coming within a certain range of events in the detection sequence^68^. These ions tend to also be detected close to each other, indicating that there are likely to originate from the same emitting site on the specimen surface^71^. These correlations offer opportunities for additional data processing steps to help identify a molecule from its detected fragments.

Here, we therefore performed APT analysis on *Sphaerechinus granularis* sea urchin spines (Figure 1a-b) and synthetic calcite crystals doped with PHNQs extracted from purple *P. lividus* spines (Figure 1c-d). We focused on the secondary spines of the *S. granularis* species because they can exhibit distinct purple– and white-colouration in separate regions (Figure 1a-c). Having these two colourations in the same biomineral is in principle a good study case to evidence the presence of PHNQ in the purple region as compared to the white one. PHNQs of purple *S. granularis* primary spines (Figure 1b) were also extracted to determine the main PHNQ molecule responsible for the purple coloration in those samples.

**Figure 1.**
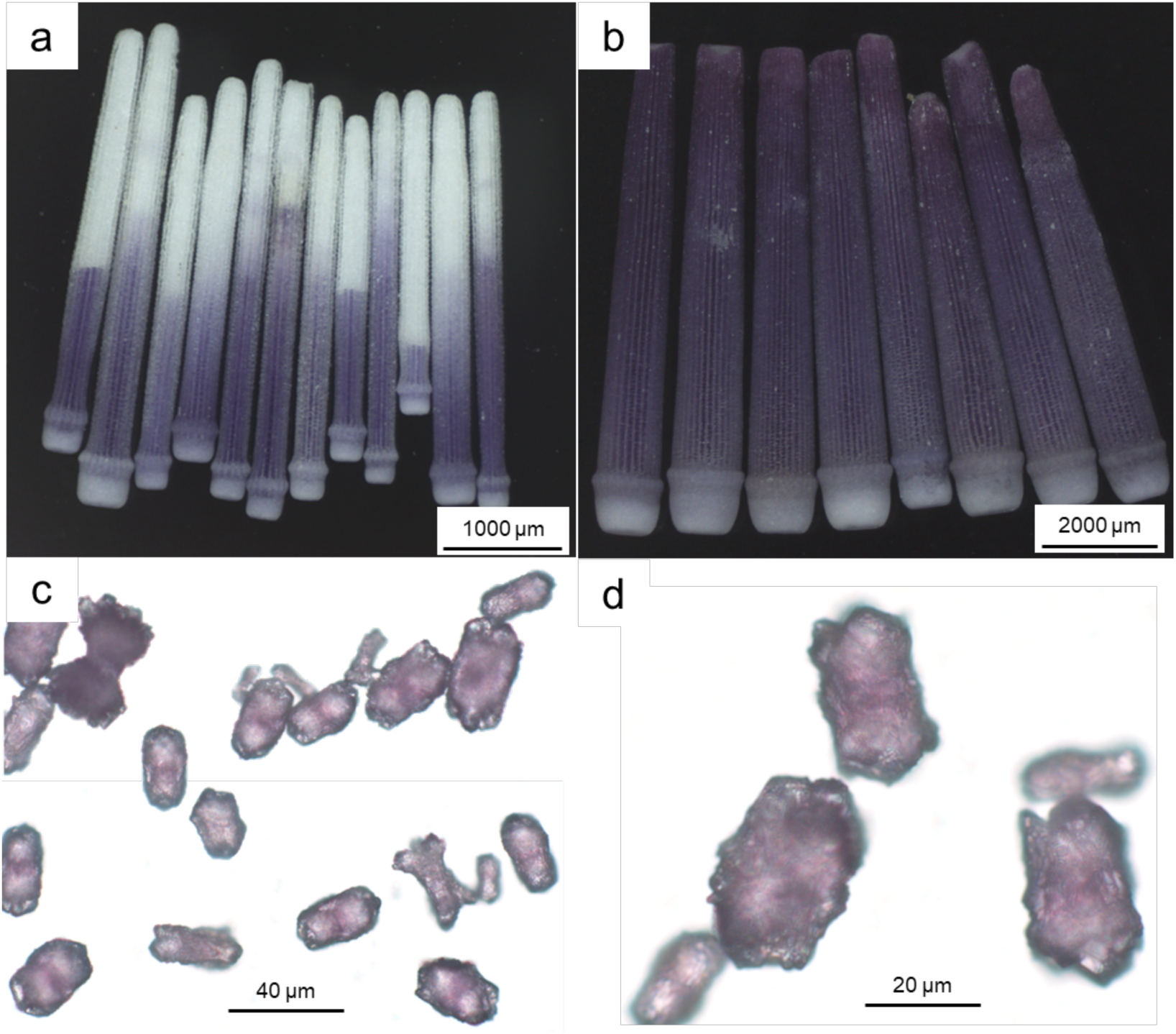
(a-b) Stereo microscope images of sea urchin spines from species S. granularis. (a) Secondary spines, which are purple-coloured at the base and white at the tip and (b) purple primary spines. (c-d) Transmission optical microscope images of synthetic calcite crystals doped with PHNQs extracted from purple-coloured spines from species Paracentrotus lividus. Doped crystals display anisotropic morphology and have purple colouration due to pigment molecules spinochrome A.

To target possible fragments related to PHNQs, we identified peaks present in purple-coloured spine regions that were absent in the white regions of the same spine. We also compared the peaks detected in synthetic pure and PHNQ doped calcite crystals (Figure 1d). Further peaks detected in both the white and purple region of the spine highlight compositional complexity with respect to pure synthetic calcite, arising from the occluded biomolecules and/or inorganic ions. To facilitate the identification of molecular fragments, we use data filtering techniques that exploit the correlation between field evaporated ions^66,72^ and have been used to analyse analytical field-ion microscopy data^68^. Proximal events containing possible PHNQ fragments indicate their fragmentation paths during field evaporation and enable chemical composition to be identified for several detected peaks. Furthermore, we can report that PHNQs have a homogeneous distribution at the nanoscale, in combination with a strong association to Mg spatial distribution, which suggests PHNQs are bound to Ca as well as Mg ions in the crystal lattice.

## 2. Methods

### 2.1 Materials

Synthetic calcite crystals with a (100) orientation was sourced from Alineason Materials Technology GmbH. Reagents were purchased from Sigma-Aldrich and used without further purification. These included calcium chloride dihydrate (CaCl_2_.2H_2_O), 1 M sodium hydroxide (NaOH) solution, ammonium carbonate ((NH_4_)_2_CO_3_), absolute ethanol, sodium hypochlorite (NaClO), glacial acetic acid, diethyl ether and sodium carbonate (Na_2_CO_3_). A Millipore Milli-Q® Integral 10 water purification system was used to obtain double distilled water (DDW) with a resistivity of 18.2 MΩ.cm at 25 °C and TOC < 5 ppb. All solutions were prepared in DDW.

*Paracentrotus lividus* and *Sphaerechinus granularis* sea urchins were obtained from the Roscoff biological station of Sorbonne University (France). Secondary spines from *S. granularis* were resin embedded using Epofix and hardener at room temperature. A cross-section through the spine was exposed by grinding up to 4000 grit and diamond polished up to 1 µm using Si paste before being washed thoroughly with milliQ H_2_O. Spines were then coated with a conductive 10 nm carbon layer. SEM images were taken with the backscatter detector in a FEI Helios 600i SEM/FIB with an acceleration voltage of 5 kV and current of 86 pA.

Polyhydroxylated naphthoquinone molecules (PHNQs) were extracted from purple *Sphaerechinus granularis* and *Paracentrotus lividus* sea urchin primary spines following^11^. Spines were treated with 5% NaClO for 8h to remove soft tissues and again for 4h after powdered (<200 μm). The spine powder was slowly dissolved with 10% acetic acid. The resulting solution was centrifuged to remove the organic acid-insoluble matrix (AIM). The supernatant containing the acid-soluble matrix (ASM) was filtered through a 1 kDa membrane. The ASM fraction below 1 kDa (ASM<1 kDa), containing free PHNQs, was extracted with equal volume of diethyl ether after having adjusted the pH to 2. The diethyl ether phase containing the PHNQs fraction was recovered, evaporated and further freeze dried. FTIR spectrum of ASM < 1 kDa was recorded using a Spectrum 400 FT-IR/FT-NIR Spectrometer (PerkinElmer) with a Universal ATR sampling accessory. Four scans were performed from 550 to 4000 cm^−1^ with a 4 cm^−1^ step resolution. HR-MS analysis was performed using a LTQ Orbitrap XL spectrometer with ESI in negative. ASM < 1 kDa was solubilized in 100 μL methanol. m/z values were determined with a set resolution of 30,000.

Doped calcite crystals were precipitated by the ammonium carbonate vapor-diffusion method in the presence of PHNQs using a 16-well Nunc™ Lab-Tek™ chamber slide system. Here, wells were filled with 400 µL of a solution containing 10 mM CaCl_2_, 40 mM NaOH and 0-50 µg mL⁻¹ PHNQs. The loaded 16-well slide was placed on the desiccator’s perforated porcelain plate, above a Petri dish containing 2.5 g of solid (NH_4_)_2_CO_3_. The Petri dish and wells were covered with Parafilm and punctured with a needle, allowing controlled diffusion of CO_2_ and NH_3_ vapor into the wells. The reaction proceeded undisturbed for 3 days at room temperature in a chemical hood, after which the supernatant was carefully removed and the residual solution aspirated with a suction pump. The precipitated crystals at the base of each well were rinsed sequentially with deionized water and absolute ethanol to remove soluble salts and unincorporated pigment, then dried under vacuum prior to further characterization.

Stereo microscopy images were obtained using a Zeiss Stemi 508 equipped with an AxioCam ICc 5 camera, with a magnification ratio of 0.63 to 2. Samples were illuminated with a polarized ring light and images were acquired with the Zen core v3.8 software. Light microscopy images were obtained using a Zeiss AxioImager A2 POL, equipped with an AxioCam CCD camera. Images were acquired with the Zen core v2.7 software.

### 2.2 APT specimen preparation and analyses

Series of needle-shaped specimens for APT were prepared by a site-specific lift out using the methodology introduced by Thompson et al.^106^ using a scanning-electron microscope / focused-ion beam (SEM/FIB). A commercial Si-based coupon (Cameca Instruments Ltd, Madison, WI, USA) was used as support. APT analyses were performed in a Cameca LEAP 5000 XR in laser pulsing mode, at a wavelength of 355 nm, with a laser pulse energy of 120–150 pJ and a laser pulse frequency of 100 kHz for all analyses. The base temperature of the specimen was maintained at 50 K, and the average detection rate fixed at 5 ions per 1000 pulses. Note that all the mass spectra displayed were calculated using a quadratically increasing bin size, which explains the flat noise level at lower masses.

### 2.3 APT evaporation field estimation

To verify that the three datasets were acquired under comparable field conditions, we used ratios of detected known elements at different charge states^107^. It should be acknowledged that this approach has limitations, particularly for non-conductors^108^, as many of the detected ions can arise from the fragmentation of molecular ions not considered in the model introduced by Kingham et al.^107,109^. If the majority species in the mass spectrum is Ca^2+^ ions at 20 Da. However, the series of low peaks starting at 40 Da are attributed to MgO^+^ rather than Ca^+^ (see Figure S 3 and Table S 2), making it impossible to use the Ca charge-state ratio to estimate the electrostatic field conditions. Given the lack of these and other suitable combinations of charge states of a single molecular ion, we settled on applying the fraction O_2_^+^/(O^+^+O_2_^+^) as a proxy for the electric field (as introduced by Gordon et al). Analysis across three oxides indicates that it provides a reasonable estimation even if the trend is not fully monotonous as in other reports^29,108^. The observed discrepancies may be due to alterations in the fragmentation path leading to the detection of these different ions^69,110^.

## 3. Results & Discussion

### 3.1 APT analyses of synthetic calcite crystals

A mass spectrum from the APT analysis of synthetic calcite crystals is plotted in Figure 2a. We identify most peaks (shaded in blue) as combinations of Ca, O and C, consistent with what has been reported for biogenic, synthetic and geological calcite minerals^73–75^. For example, the peaks at 20, 40, 44 and 56 Da are respectively attributed to Ca^2+^, Ca^+^, CO_2_^+^ and CaO^+^. Similar non-stoichiometric composition (10 at. % C, 34.5 at. % O) was observed due to preferential losses of C and O ions compared to Ca ions. This can be rationalised by the strongly delayed field evaporation of Ca, particularly noticeable in the high level of background following the Ca^2+^ peak at 20 Da, which can obscure C-containing peaks as observed in β-tricalcium phosphates^76^ and discussed by Schwarz et al.^32^. Further ions may also remain undetected due to the desorption of neutral molecules like CO and CO_2_^77,78^. These discrepancies can be ignored here as our goal is not to provide a quantitative analysis of calcite, known to be challenging for these materials^74,79^, but rather use as a reference baseline for the peaks detected in the PHNQ doped calcite crystals and the biomineral spine.

**Figure 2.**
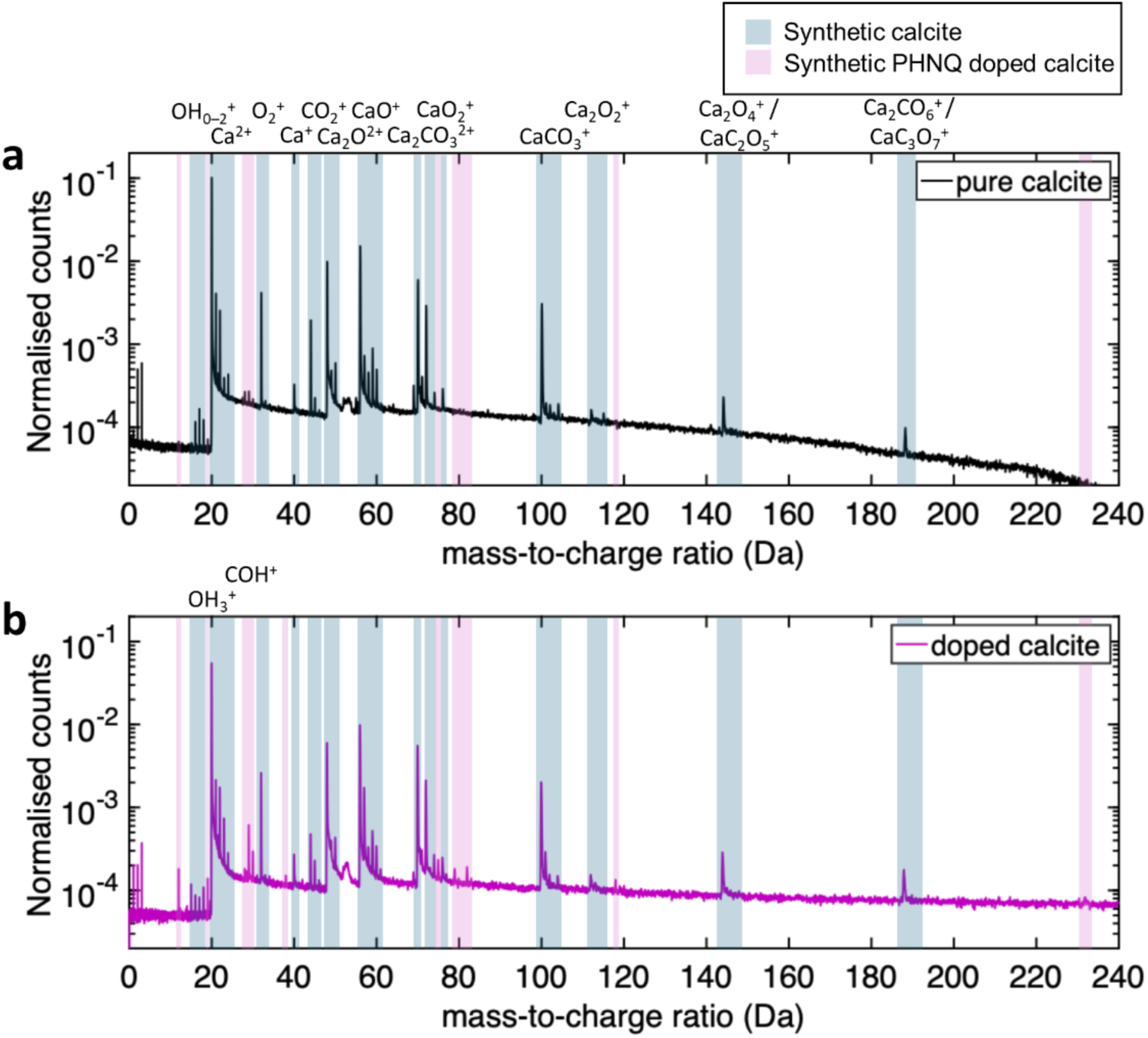
Mass spectrum in the range 0–240 Da for (a) the synthetic calcite single crystal and (b) the calcite crystal doped with spinochrome A, each normalised to the total number of ions in the mass spectrum. The ranges of mass-to-charge state ratios shaded in blue correspond to sets of peaks detected in the pure calcite and doped calcite while those in pink are peaks found primarily in the doped calcite or those with a noticeable change in intensity between the two datasets.

### 3.2 APT analyses of synthetic calcite crystals doped with PHNQs

PHNQs were first extracted from purple *P. lividus* spines (see Experimental section) that contain predominantly spinochrome A, as previously identified by ^1^H and ^13^C NMR as well as HR-MS^11^. The FT-IR spectrum of the PHNQ extract used in this study shows the typical bands of spinochromes as well as some bands attributed to impurities originating from fragments (<1KDa) of proteins, lipids and polysaccharides (Figure S 1a). Second, calcite crystals were synthesised by vapour diffusion of (NH_4_)_2_CO_3_ into a 10 mM CaCl_2_ solution^5,80^ containing 0 or 50 µg mL⁻¹ of PHNQs (see Experimental section). The crystals doped with PHNQs have a purple-pink colour and exhibit anisotropic shapes and crystallographic facets (Figure 1c-d). They are also smaller than the pure calcite crystals that are transparent and show typical rhombohedral shapes (Figure S 2). The purple coloration is homogeneous across the doped crystals suggesting that spinochrome A is evenly distributed throughout the calcite mineral at the microscale.

A mass spectrum from the APT analysis of a synthetic doped calcite crystal is plotted in Figure 2b. It shows a multitude of peaks, whose positions are reported in SI (Table S 1). Some of the peaks are the same than in the mass spectrum of calcite and are highlighted in blue, while additional peaks are marked in pink when they appear in the doped calcite and are either absent – for instance, at 79, 82 and 232 Da – or have a very different amplitude in calcite, including 12, 19, 29 Da. Electrostatic field conditions are known to influence compositional analysis^81^ as well as fragmentation of biomolecules in APT^82^. They can be evaluated based on the charge state fraction of O_2_^+^/(O_2_^+^+O^+^) across the different datasets (see Methods). Here, the charge state fractions for both datasets were between 0.93–0.94, implying comparable electrostatic field conditions throughout the measurement of both samples. Therefore, the additional peaks as well as the peaks having a higher intensity in the doped calcite can be attributed to fragments from the additional organic species, in this case spinochrome A, although some organic impurities might also be present in minority. Peaks at 19 and 29 Da are commonly found in the analysis of the organic regions of mineralised bone^32^, and considered as H_3_O^+^ and COH^+^ respectively, as indicated in Figure 2b. The peak at 29 Da is present in relatively higher concentration in the doped calcite and could correspond to the fragment 7 or 12 or labelled A (COH^+^) in Figure 3a and 3b, respectively. The peak at 12 Da is most likely C^+^ (fragments 3, 4, 9).

**Figure 3.**
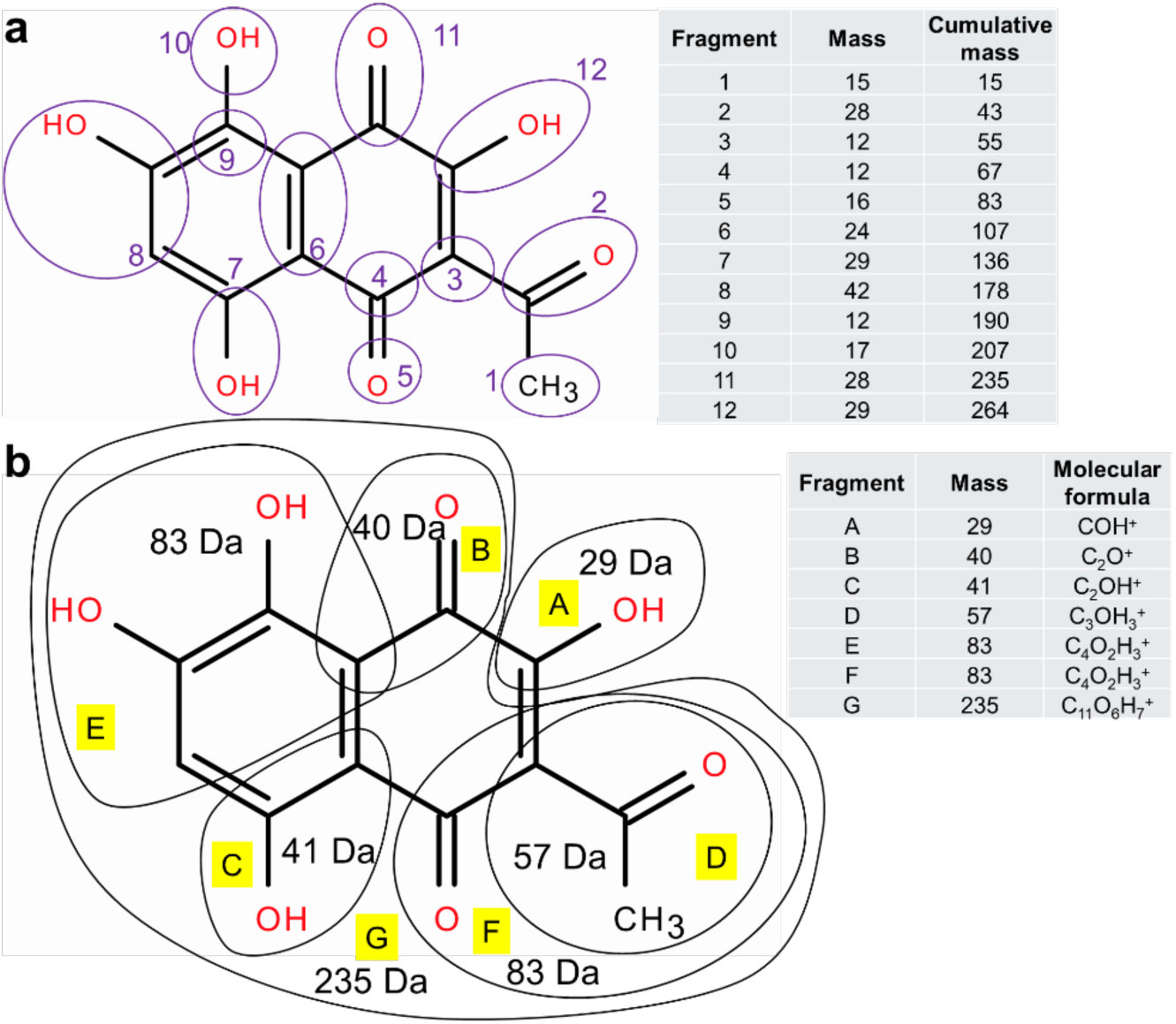
(a) Molecule of spinochrome A (drawn using ChemDraw) with potential fragments with their respective mass reported in the table, along with the cumulative mass from adding fragments together. (b) Other possible fragments with masses corresponding to some found in the mass spectrum of the doped calcite. These are each listed alongside in the table with corresponding mass and molecular formula.

The peak at 40 Da has a similar intensity in both pure and doped synthetic calcite, suggesting it is Ca^+^ and not the fragment marked B (C_2_O^+^) in Figure 3b. No peak at 41 Da could be detected that could correspond to fragment C (C_2_OH^+^) in Figure 3b. The peak at 44 Da corresponding to CO_2_^+^ can originate from both organic molecules and calcite. In this case, it is more likely to come from the calcite itself, as it appears at similar intensity in both pure and doped synthetic calcite. The ratio of the amplitude of peaks 44 and 45 Da (HCO_2_^+^) are 0.22 in the pure calcite and 0.60 in the doped calcite, indicating a higher probability of CO_2_ molecules becoming protonated in the doped calcite by organic PHNQ molecules during molecular ion formation, given that field conditions are comparable. The peak at 82 Da could correspond to the fragments E or F (C_4_O_2_H_3_^+^) after the loss of a proton (Figure 3b). The peak at 232 Da likely corresponds to the fragment G (C_11_O_6_H_7_^+^) (235 Da) in its deprotonated form. This fragment nearly corresponds to the complete spinochrome A molecule having lost an COH^+^ group (29 Da) and assuming deprotonation. Deprotonation of hydroxy groups in spinochrome A is responsible for its purple colouration in solution at pH>7 and is proposed to occur during the formation of both synthetic and biogenic calcite before being occluded within the crystals^11^.

Deprotonation of molecules can also occur during field evaporation in APT analysis^53,83^, as reported for glucose molecules^41^. Further peaks were identified at 75 and 79 Da in the doped calcite. The peak at 75 Da may be glycine, which could originate from organic impurities in the PHNQ extract or protonated diethyl ether which is used as a solvent for extraction. The assignment to glycine appears more likely as it is the main amino acid found in the proteins of the acid-soluble matrix extracted from *P. lividus*^5,8^. The identity of the peak at 79 Da is unknown, but could be related to benzene, which would have a mass of 78 Da, however further evidence would be required to confirm this interpretation.

### 3.3 APT analysis of *S. granularis* secondary sea urchin spine

Prior to the APT analysis of *S. granularis* secondary sea urchin spines, which exhibit a white tip and a purple base (Figure 1a), PHNQs from the purple primary spines were extracted and analysed by HR-MS to determine their composition (see Methods). The HR-MS spectrum of this PHNQ extract shows the major presence of spinochrome A (>95%) with the minor one of spinochrome B and C (Figure S 1b). These results indicate that spinochrome A is involved in the purple colour of the *S. granularis* species as it is for the *P. lividus* one^11,62^.

Figure 4a-c display stereo microscope images of the polished cross-section of a *S. granularis* secondary spine after the preparation of APT specimens. As for the doped calcite crystals, the purple region at the base of the secondary spine shows a homogeneous coloration at the microscale (Figure 4c). APT specimens were obtained from the purple and white regions of this spine in the areas targeted in the scanning-electron microscope equipped with a focused-ion beam (SEM-FIB)^84^ (see Methods) and indicated by the black boxes in Figure 4d-e.

**Figure 4.**
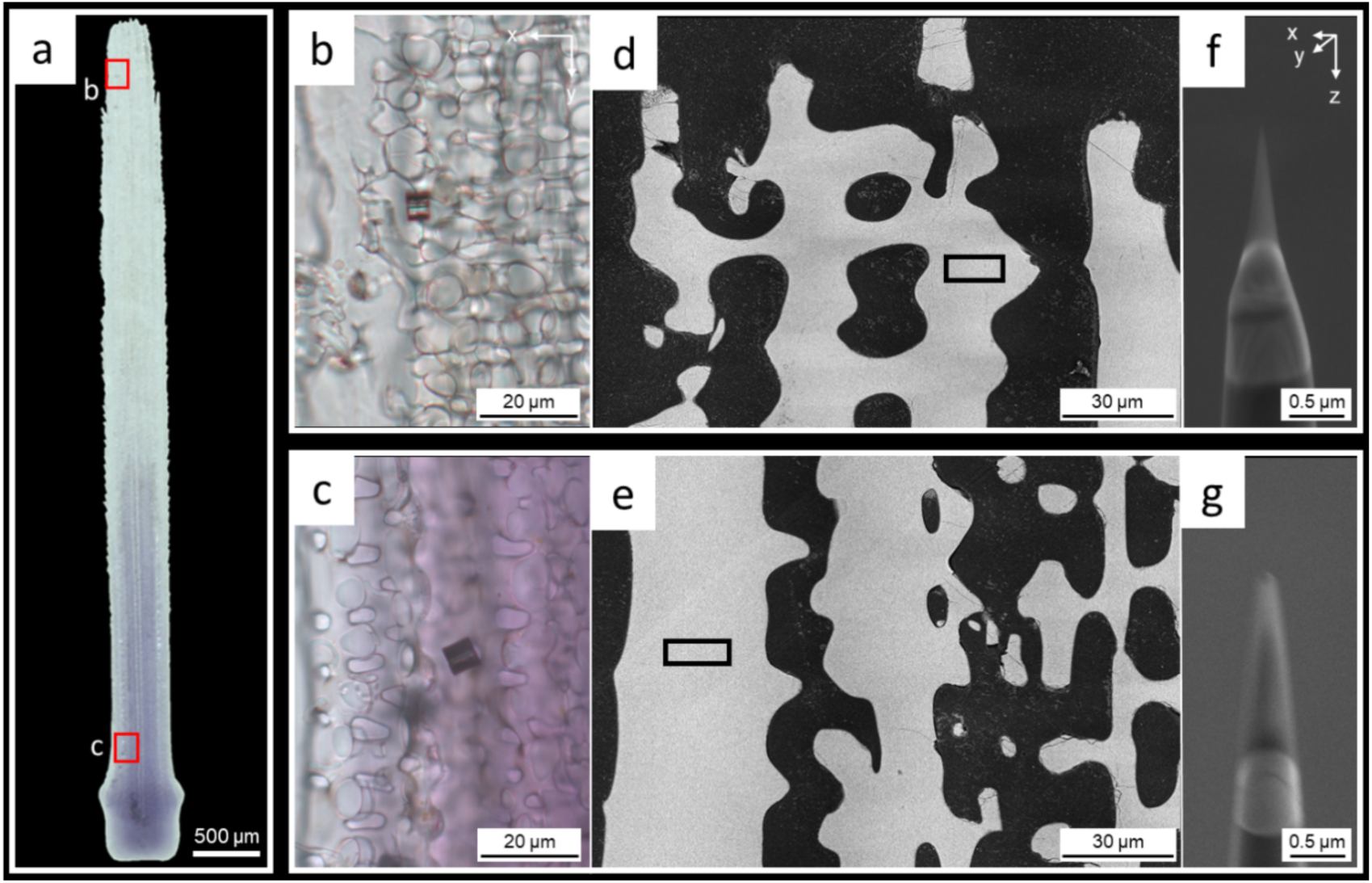
(a-c) Stereo microscope images of the polished cross-section of an embedded secondary spine from Sphaerechinus granularis after APT specimen preparation, which has a white tip (b) and purple base (c) and (d-e) SEM images taken in BSE mode of the white and purple regions respectively before the preparation of APT specimens and of the APT specimens themselves. The black boxes indicate the location of the APT specimen shown in f and g.

The mass spectra obtained from APT analyses of the white and purple regions of the spine are plotted in Figure 5a and 5b, respectively. The O_2_^+^/(O_2_^+^+O^+^) charge state fraction for both regions ranged between 0.97–0.98. As this value is within a few percent of 0.93–0.94 recorded for synthetic pure and doped calcite crystals, the analysed datasets were acquired under comparable electrostatic field conditions. We therefore consider that the calcite mineral phase of all samples will fragment similarly and that differences in peaks detected are due to compositional variation. It should also be noted that organic molecules, including spinochrome A, will likely have a different bonding environment in synthetic and biogenic calcite, which could also affect the calcite phase itself, impacting fragmentation patterns and dictate the ions that are detected.

**Figure 5.**
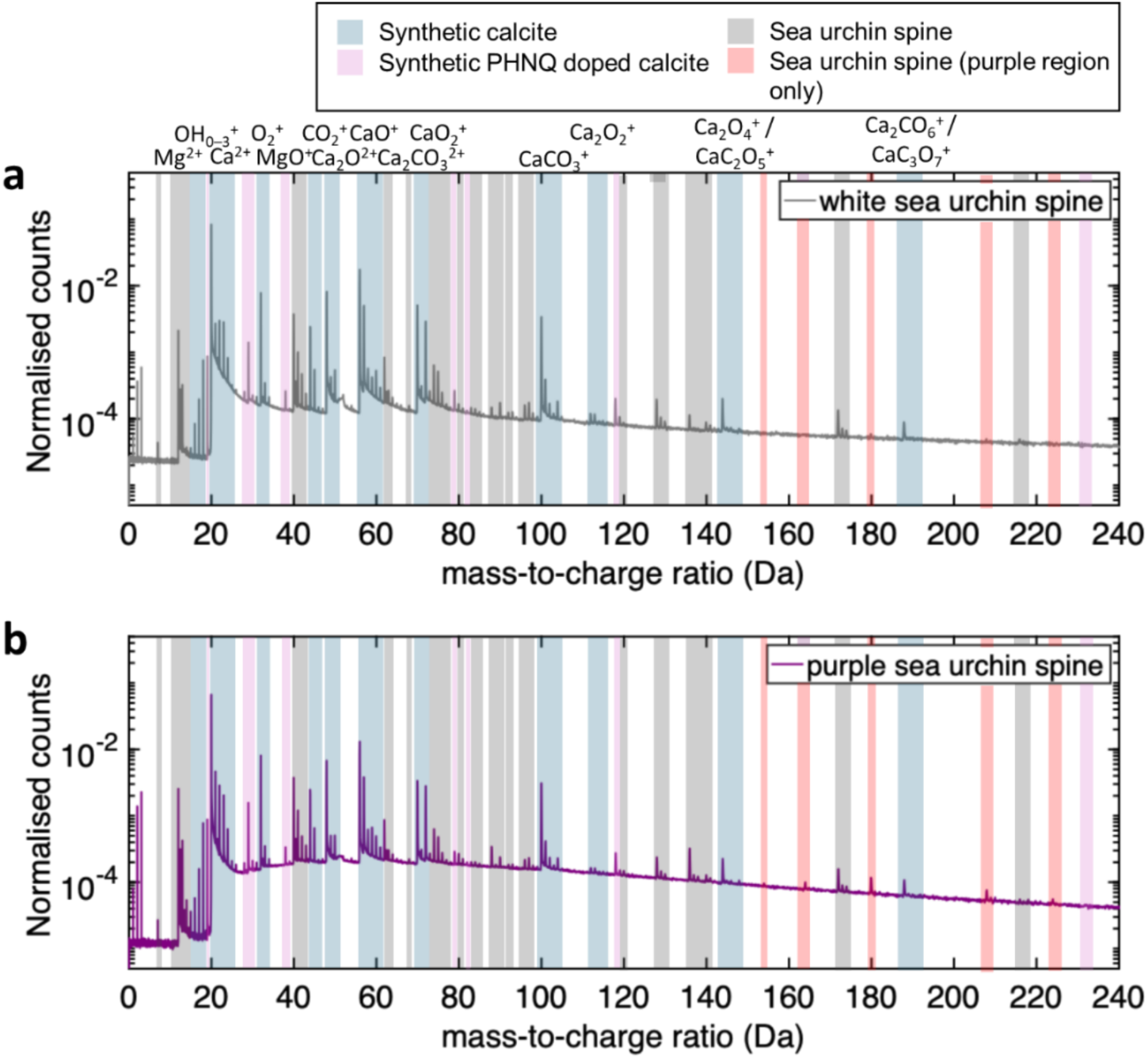
0 – 240 Da section of the mass spectra of the white and purple regions of the sea urchin spine, each normalised to the total number of ions in the mass spectrum. As for Figure 1, peaks detected in the calcite single crystal are shaded in blue and peaks found primarily in the doped calcite or those with a noticeable different intensity between the two datasets are shaded in pink. In addition, peaks appearing only in the white and urple regions of the spine are marked in grey and peaks specific to the purple region are marked in red.

The positions of the peaks identified above for the synthetic pure and doped calcite are indicated in blue and pink respectively in Figure 5. Peaks detected in both the white and purple regions of the sea urchin spine, but absent in the synthetic samples, are marked in grey, whereas additional peaks detected only in the purple region are marked in red. Thus, peaks marked in grey provide valuable information on the nature of the organic molecules and inorganic ions comprising the biomineral while those marked in red are most likely to originate mainly from spinochrome A. Several peaks including those at 19, 29, 40 and 57 Da are present in both synthetic and biogenic calcite, but exhibit notably higher intensities in the spines. In the synthetic pure and doped calcite crystals, these peaks were attributed respectively to OH_3_^+^, COH^+^, Ca^+^ and CaO^+^ with some organic fragments. Some of these peaks, for example at 19 and 29 Da, may be generic fragments of organic molecules, possibly representing hydroxyl and carbonyl groups respectively, and are therefore marked with grey backgrounds in Figure 4a-b. However, the peak at 40 Da is here attributed to MgO^+^ rather than Ca^+^ due to its higher intensity in biogenic calcite with respect to synthetic ones, and also due to the corresponding isotope peaks at 41 and 42 Da (discussed in methods). We can note that the ratio of Mg to Ca derived from the analysis of these mass spectra is 5.6 % in the white region and 7.9 % in the purple region of the spine. This corresponds to a Mg concentration of 2.1 at. % in the purple and 1.8 at. % in the white, however this could be overestimated due to losses of C and O. These values appear to fall in range for those already reported for *S. granularis* spines^85^. In the spine, the peak at 57 Da may correspond to a common organic fragment, for example a glycine molecule with the loss of a OH group (C_2_ONH_3_^+^). As this peak also appears in the synthetic doped calcite at a higher intensity than pure calcite, it could also correspond to C_3_OH_3_^+^, marked fragment D in Figure 3b. It should be noted that peaks are also present in the sea urchin spine at 23 Da, likely corresponding to Na^+^, which were also detected in synthetic calcite, albeit at lower intensity.

In more details, peaks found between 0 and 19.5 Da, plotted for both spine regions in Figure 6a-b, are attributed to Mg^2+^ and NH^+^/CH_3_^+^, based on their *m/z* values and due to their absence in the mass spectrum of synthetic pure and doped crystals, with the latter plotted in Figure 6c.

**Figure 6.**
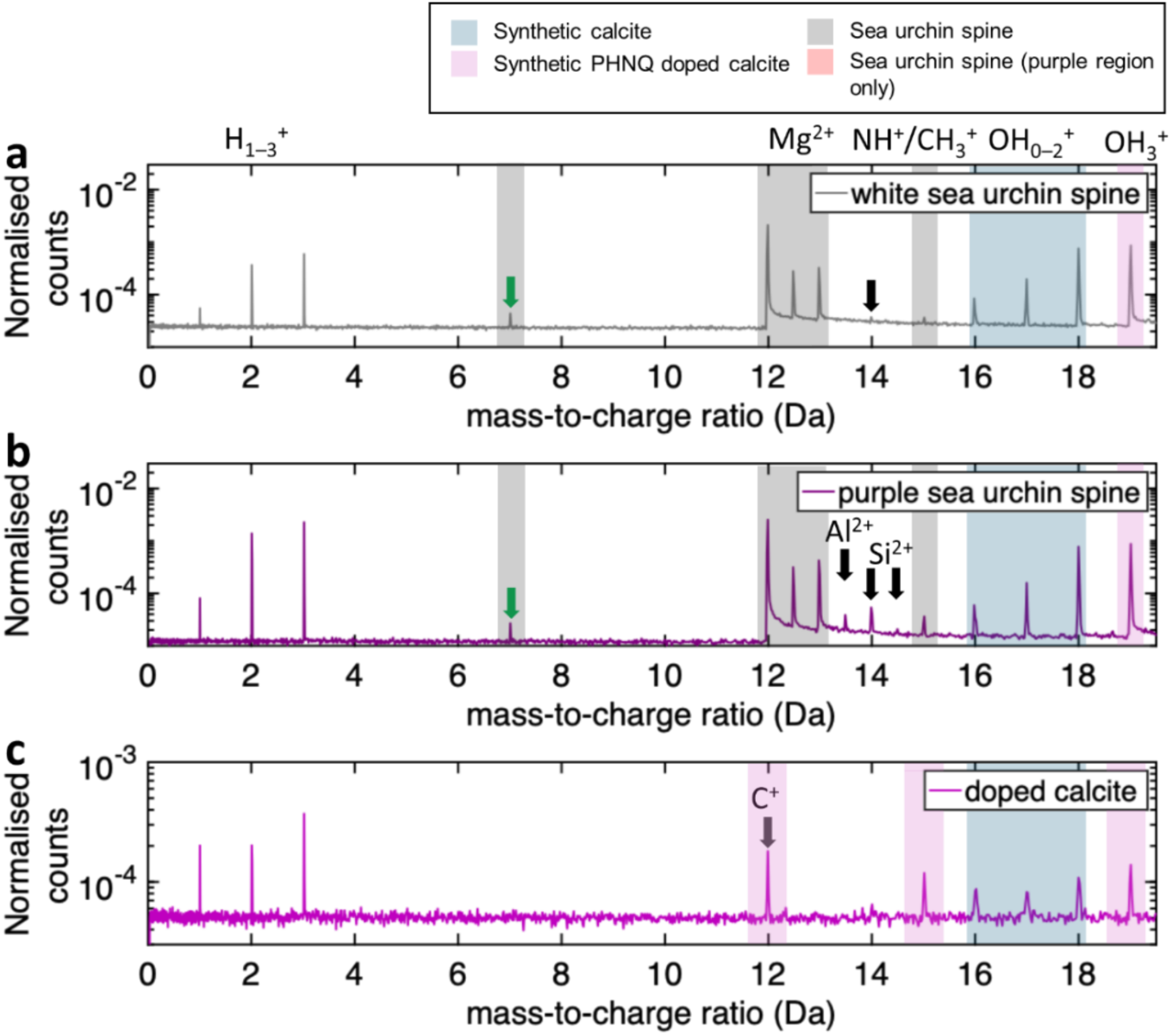
Sections of the mass spectra for (a) the white region, (b) the purple region of the sea urchin spine and (c) the doped calcite between 0 and 19.5 Da. All spectra are normalised to the total count in the dataset. Note that the y-scale was adjusted between (a–b) and (c). Sets of peaks corresponding to a range of ions appearing in spine regions are specifically labelled in grey in (a-b), and marked in blue when also found in synthetic calcite. Peaks detected at 7, 13.5 and 14-14.5 Da are highlighted by arrows and identified as Li^+^, Al^2+^ and Si^2+^ respectively. In (c), the pink background corresponds to peaks belonging exclusively to spinochrome A.

The three spectra show peaks at 19 Da corresponding to OH_3_^+^, yet with the doped calcite being an order of magnitude lower than the biomineral (the y-scale was adjusted for visibility). The OH_3_^+^ and NH^+^/CH_3_^+^ peaks are often detected in the analysis of protein molecules^44,45,86^, suggesting these belong to organic molecules contained within the spine. Mg is also a known component of sea urchin spines^85,87^ and is studied in more detail below. In the purple region of the spine, there are additional peaks that can be assigned to Al^2+^ and Si^2+^ – the latter could be attributed to N^+^; however, an additional peak at 14.5 Da, corresponding to ^29^Si^2+^, exhibits an isotopic ratio close to the natural abundance (0.065 vs. 0.051) and therefore favours the Si^2+^ assignment. The presence of Si and Al in the purple region could indicate they are bound to spinochrome molecules, similar to the relationship between echinochrome A and metal ions including Fe^88,89^. In addition, a peak at 7 Da marked by the green arrow in Figure 6 is detected in both regions but is absent in the synthetic crystals. The assignment of this elemental identity is discussed in detail below.

Peaks between 72 and 240 Da are plotted for the white and purple regions of the sea urchin spine and compared with those of the doped calcite crystal in Figure 7. Here, several additional peaks (shaded in grey) are present in the mass spectra of the spine while they are absent in the synthetic crystals. These include peaks at 128, 136, 140 and 172 Da that can be reasonably assumed to originate from fragments of proteins and saccharides within the biogenic calcite^5,8,18,59^. In particular, sea urchin pigmented spines are known to comprise ASM proteins that include the well-studied basic non-glycosylated protein SM50 and acidic glycoprotein SM30, which are mainly composed of glycine (75 Da) and aspartic acid (133 Da)^18,59^. Given that these listed peaks are all followed by lower intensity peaks at +1 and +2 Da with comparable intensity to the abundance of Mg isotopes, it is possible they comprise a combination of fragments of larger organic molecules and Mg ions.

**Figure 7.**
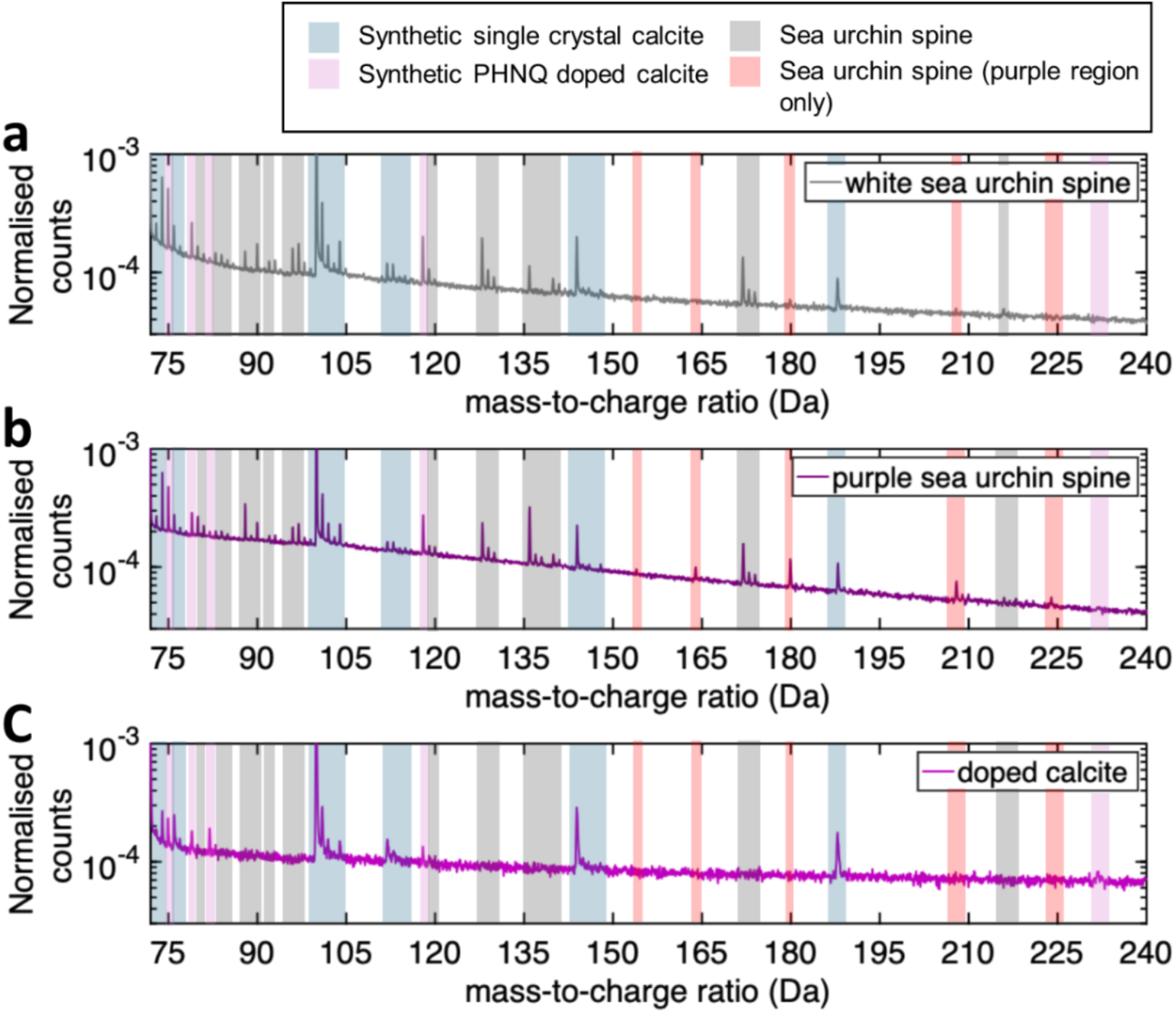
72–240 Da section of the mass spectra of the (a) white and (b) purple regions of the sea urchin spine, along with (c) the doped calcite crystal. The coloured ranges are indicated by the legend, with five individual peaks which appear only in the purple spine region and not in the white are marked with arrows.

In the purple spine region, there are three main additional peaks at 164, 208 and 224 Da that do not appear in the white spine region and one with significantly higher intensity at 180 Da (highlighted in red and marked by arrow in Figure 7b). The 180 Da peak is the most intense and could correspond to glucose, which has been identified as part of the monosaccharide contents of the organic matrix in the spines of *Arbaxia lixula* together with glucosamine and galactosamine (C_6_H_13_NO_5_) (179 Da)^10^. Complete protonated glucose molecules^41^ were detected at 181 Da by APT analysis of a saturated glucose solution, alongside lower mass fragments, including at 163 Da. Other molecules such as hydroxyproline (C_5_H_9_NO_3_) or glutamic acid (C_5_H_9_NO_4_)^32^ were proposed to be detected in mouse femur based on the molecular ions measured by APT. These studies suggest that some organic molecules may resist fragmentation under an electric field and enable larger molecules to be detected. The peak at 164 Da could then come after the fragmentation of glucose removes one O atom. However, organic fragments COH^+^ and OH_3_^+^ have comparable intensities in the white and purple regions (Figure 5a-b) suggesting organic matrix composition does not vary enough to explain additional peaks in the purple region. Therefore elevated 180 Da peaks in the purple region could result from additional monosaccharide fragments originating from macromolecules known to be complexed with spinochrome A in purple *P. lividus* spines^11^, or be direct fragments of spinochrome A molecules.

The peak at 208 Da is highest in intensity after 180 Da. This peak would therefore be the most likely candidate to be a fragment of spinochrome A, particularly as this ion would constitute a full molecule which has lost two CO^+^ groups (C_10_H_8_O_5_^+^). It is possible, therefore, that this fragment could lose a further CO^+^ group to give a peak at 180 Da (C_9_H_8_O_5_^+^), or CO_2_ to give the peak at 164 Da (C_9_H_8_O_4_^+^). If it is assumed that spinochrome A is deprotonated, the peak at 180 Da could also directly be a fragment of spinochrome A without the ion outlined as fragment D (mass 82 Da) in Figure 3b, supporting the detection of a peak at 82 Da in the doped calcite (Figure 2b). The peak at 224 Da represents a loss of 40 Da from spinochrome A, which is fragment B in Figure 3b. Alternatively, assuming again deprotonation of spinochrome A, the peak could originate from the loss of a fragment with a mass of 38 Da, which is present in the doped calcite (Figure 2b) and is possibly a combination of two molecules of the common organic fragment OH_3_^+^ (O_2_H_6_^+^). It could additionally be possible that the peak at 224 Da is from spinochrome C molecule (original mass 279 Da – Figure S 1b) which has lost two CO groups. Altogether, as it is very challenging to assign these peaks to organic fragments, we require additional data processing techniques which can identify complex fragmentation patterns and the origin of specific molecules.

Therefore, to facilitate the identification of these peaks, a correlation histogram^90^ was calculated and is plotted within two different ranges of mass-to-charge state ratios in Figure 8a and 8b. Such histograms highlight specific combinations of ionic masses detected on multiple events (i.e. when more than a single ion was detected following a single laser pulse). The level of background in the histogram is high, in part originating from the Ca-containing ionic species, as expected from previous work on Ca phosphate^76^. The presence of long tails extending from 0 Da suggests continuous field evaporation at the DC voltage, uncorrelated to the laser pulse. No strong correlations can be found between high mass fragments to suggest they originate from the same molecule, specifically those seen only in the purple sea urchin. However, these long tails and high background could obscure signals.

**Figure 8.**
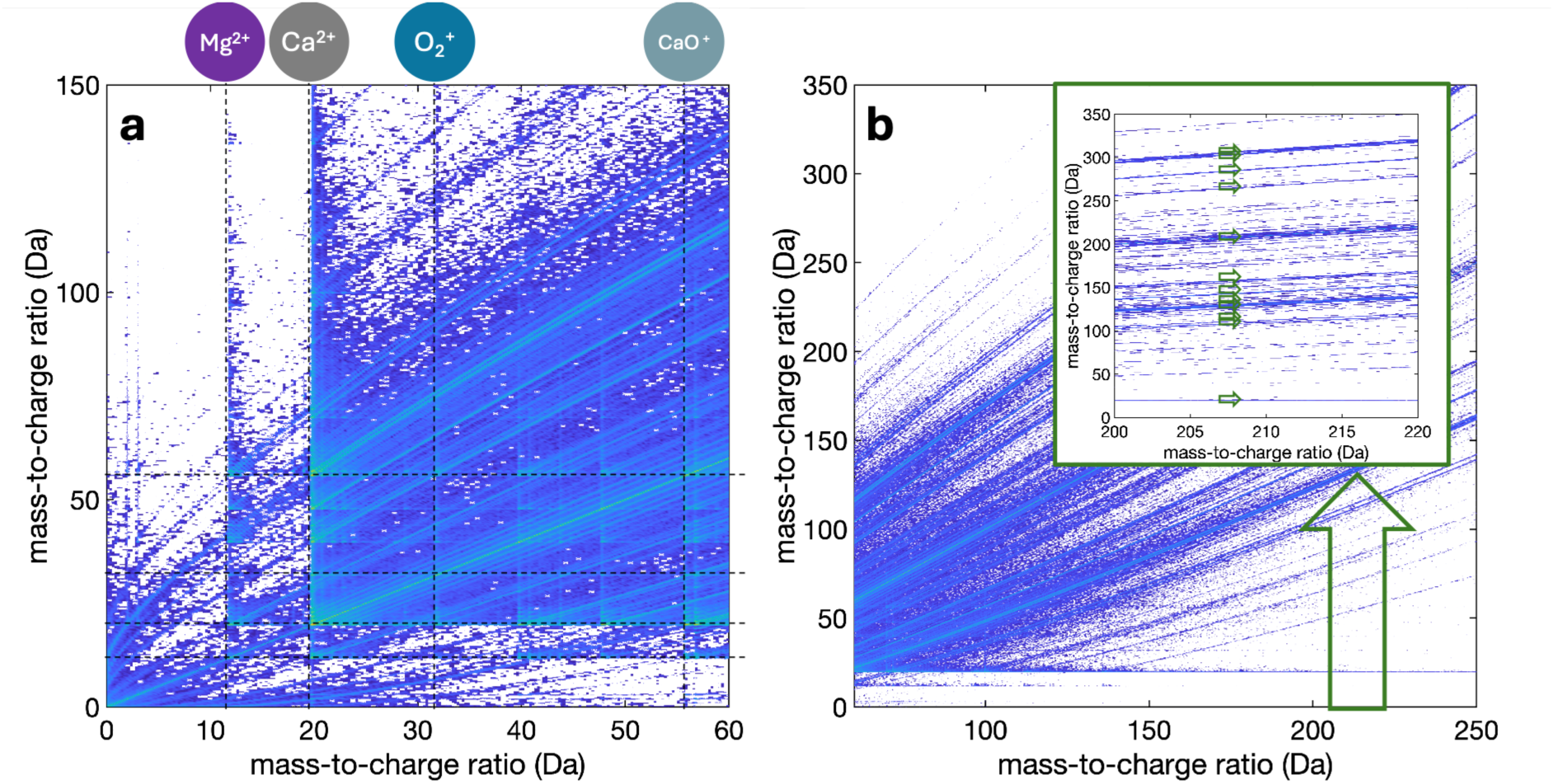
Correlation histogram for the range (a) 0–60 Da and (b) 60–240 Da for the analysis of the purple sea urchin spine. Inset is a close-up on the 200–220 Da, with green arrows pointing to the position of the major background tracks at 208 Da.

Here, to further clarify the identity of unknown peaks, we propose filtering the list of ions in the dataset based on the sequence of detection and the distance between ion impact. This approach was proposed by Morgado et al.^68^ to analytical field-ion microscopy and is akin to the methodology introduced by Saxey, but focuses on a targeted range of masses^90^. Here, it can be used to list ions with an arrival correlated with other ions from that specific mass range. Here we extracted a first list that contains the detected ions in multiple events. A second subset list events collected within a defined radius on the detector and within a certain window of number of detected events before and after ions in the targeted range. For this approach, we adopt the term *coincidence mass spectrometry*, which has been used previously for other time-of-flight based mass spectrometry approaches^91,92^. Here, we used a cut-off distance of 2 mm on the detector and ions detected within +/− 20 events based on previous work^68^ to the targeted mass range, and we refer to these as “coincidence events”.

### 3.4 Identifying the peak at 7 Da in the purple spine region

To demonstrate how information gathered from coincidence APT analysis can help identify unknown peaks, we first considered the 7 Da peak detected in the mass spectra of both regions of the sea urchin spine but not detected in synthetic calcite (Figure 6, green arrow). This peak is a good candidate for analysis due to the limited number of possible ions that could be detected within this mass range. A rapid assessment of the 7 Da peak would likely assign it to N^2+^ or CH_2_^2+^, given the presence of amino acids from proteins or other biomacromolecules embedded within the spine. However, there are several indications that suggest this may not be the most logical identification. As discussed above, the peak at 14 Da appears to be associated to ^28^Si^2+^ and not N^+^. The detection of N^2+^ without N^+^ seems unlikely, particularly due to N having a high second ionisation energy (29.60 eV)^93^. Additionally, in the white region, a peak at 14 Da is observed but with a very low intensity (Figure 6b). Furthermore, in previous reports of APT analysis on amino acids^43,44^, which are likely to be the main source of N in the spines, peaks of N^2+^ were rarely reported. Even when observed under high electric field conditions^32,82,94^, it should be consistently detected alongside C^2+^, which would also be expected from biomacromolecules. This is based on the relatively lower second ionisation energy of C compared to N (24.38 vs. 29.60 eV, respectively)^95^. Yet, herein, no peak at 6 Da is detected above the background in either the purple or white regions of the sea urchin spine. An assignment to CH_2_^2+^ is unlikely given that peaks at 6 Da and 6.5 Da are not observed, which would correspond to C^2+^ and CH^2+^, in addition to the low probability of these charge states being detected for these species.

Coincidence analysis was therefore performed for the peak at 7 Da, filtering the events that contain at least one ion in the range 6.99 – 7.02 Da in the purple region of the spine (Figure 9a). Figure 9a shows the mass spectrum for all ions detected as part of multiple events in purple, and those from the coincidence events in lilac. No notable peaks appear beyond those expected from calcite (shaded in blue). There are also no correlations with the main organic fragments identified typically at 19 Da (OH_3_^+^), 29 Da (COH^+^) and 44 Da (CO_2_^+^), suggesting that ions at 7 Da are not components of larger organic macromolecules but rather of the mineral phase. A likely candidate for alternate assignment is ^7^Li^+^, as Li_2_CO_3_ is a highly stable compound and Li substitutes for Mg in many minerals^96^. Li is also found in some marine biogenic carbonates^97^ and abundant in seawater, where its content depends on the local sea water composition^98^. The presence of Na detected here in the biominerals would support this hypothesis. An argument against assigning the peak at 7 Da to Li^+^ would be the lack of a corresponding peak at 6 Da from the other stable isotope ^6^Li^+^ with an abundance of 7.6%. The absence of the 6 Da peak in the main mass spectrum can however be explained by low ^6^Li ion counts expected based on natural abundances (^6^Li^+^ 7.6% and ^7^Li^+^ 92,4%), being obscured by the background. Furthermore, the ions in the peak at 7 Da are primarily detected as single hits (87%), hence the very low probability of a ^7^Li – ^6^Li combination in multiple events, with only ∼3 ions explaining their absence in this coincidence analysis. Together, this might explain the absence of the isotope peak at 6 Da.

**Figure 9.**
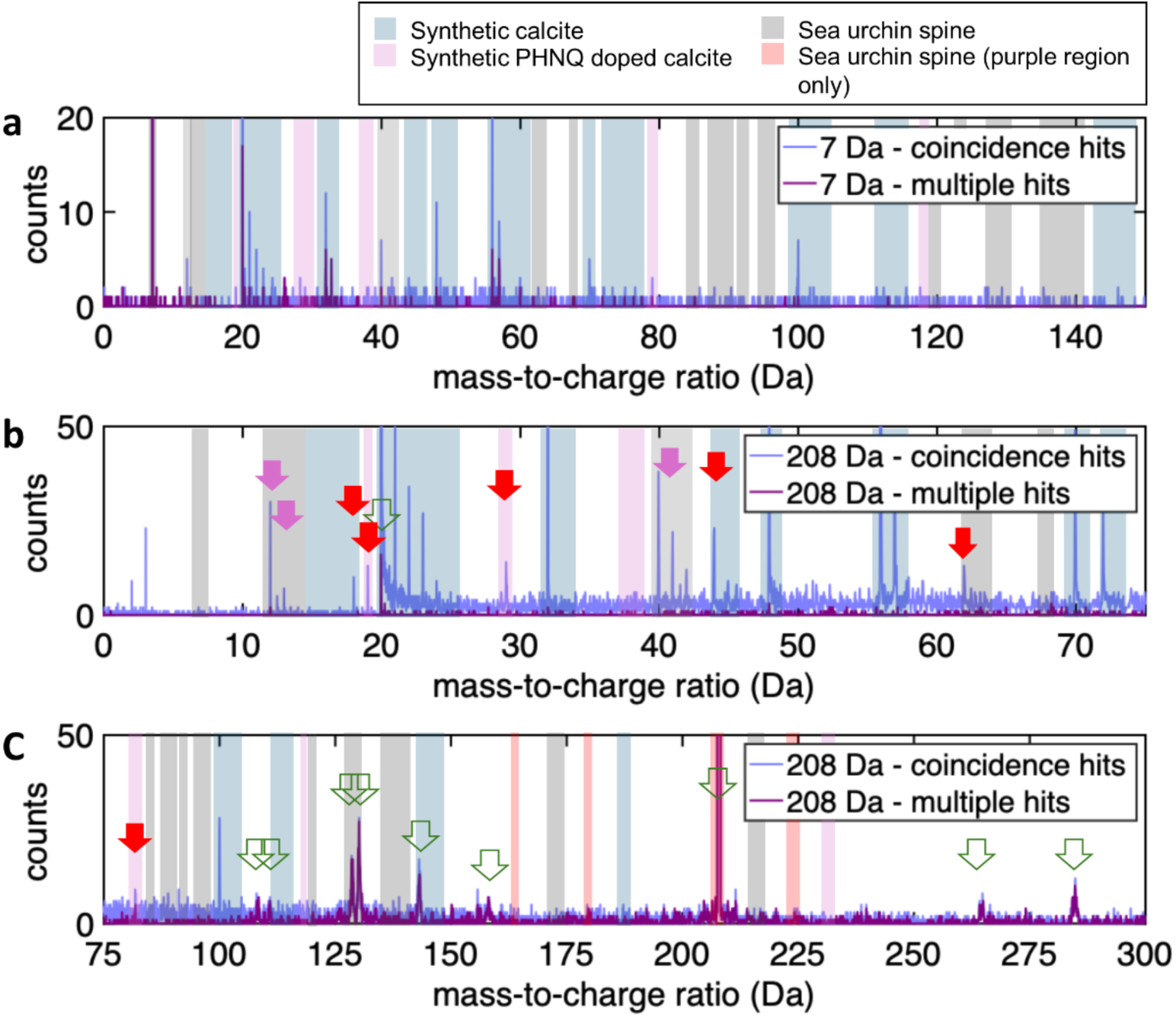
Coincidence APT for the mass spectra of the purple region of the sea urchin spine. (a) Mass spectrum for ions detected as part of multiple hits or in coincidence with those in the peak at 7 Da in the range of 0–150 Da (2mm, +/− 20 hits). (b-c) Sections of the mass spectrum for ions detected as part of multiple hits or in coincidence with those in the peak at 208 Da in the range of (b) 0–75 Da and (c) 75–300 Da. Arrows highlight peaks of interest; pink for Mg-related peaks, red for organic-related fragment peaks and green for peaks appearing from background thermal tails.

### 3.5 Identifying the peak at 208 Da in the purple spine region

A coincidence analysis was also performed for the peak at 208 Da (Figure 9b-c) as it is one of the most intense peaks that only appears in the analysis of the purple region of the sea urchin spine (Figure 7b). Its prominent appearance exclusively in the purple region makes it a good target to showcase the coincidence mass spectrometry methodology. This peak is also close in mass to the 232 Da peak observed in the doped calcite, separated only by 24 Da, indicating a difference of two C atoms and is less likely to overlap with ions from other sources, which may be the case for peaks identified at lower masses at 79 and 82 Da for example.

Two sections of the mass spectrum from 0 to 75 Da and from 75 to 300 Da are plotted in Figure 9b-c for the coincidence analysis of the 208 Da peak, filtering ions detected in either multiple or coincidence events that contain an ion in the mass range 207.8–208.4 Da. For the multiple hits, plotted in purple, numerous peaks appear alongside 208 ions at 20, 108, 111, 128, 130, 143, 158, 265 and 285 Da (indicated by the green arrows). Several of these would otherwise be below the background in the original mass spectrum. Figure 8b shows that these peaks are not from the calcite biomineral but rather formed from the tails of the peaks (marked also with green arrows) arising from the late field evaporation of calcium ions or calcium-containing molecular ions as previously reported^76^. Therefore, these peaks cannot be interpreted as more than background, and this is supported by the mass spectrum plotted separately for multiple events in Figure S 4, in which no peak appears at 208 Da.

At lower masses ranges however, the coincidence analysis of 208 Da ions contains many peaks at predicted mass ranges (Figure 9b). While most of these peaks are expected from calcite (marked by the blue lines), several ones (marked by pink arrows) at 12, 13, 40 and 41 Da likely correspond to Mg^2+^ and MgO^+^ (as discussed above). Although alternative assignments to C_2_O^+^ (40 Da) and C_2_OH^+^ (41 Da) cannot be fully discounted, the appearance of Mg-related peak in 208 Da coincidence events indicates some possible complexation between PHNQ and Mg ions. Further peaks at 18, 19, 29, 44, 57, 62 and 79 Da indicated by red arrows can be here attributed to organic fragments, several already identified as OH_3_^+^, COH^+^ or CO_2_^+^. The peak at 57 has also been discussed as an organic fragment, either directly from the spinochrome A molecule (fragment D Figure 3b), or as a protonated combination of two CO^+^ groups (C_2_O_2_H^+^). Peaks at 62 and 79 Da can be identified as organic fragments, either from spinochrome A molecules or components of organic molecules which form larger complexes with pigments. The peak at 79 Da was also detected in the doped calcite crystal. A key difference is that the peak at 232 Da observed in the doped calcite is not detected in the purple spine region in the original nor in the coincidence mass spectrum. This difference of fragmentation could be explained by the different chemical environment of PHNQs in the sea urchin spine compared to synthetic doped crystals, possibly due to the complexation with Mg or macromolecules for instance, which could form larger complexes as suggested above.

Finally, coincidence analysis for peaks at 164, 180 and 224 Da found in the purple region was also performed, see Figure S 6. Similar peaks to the 208 Da coincidence analysis were revealed, including calcite and Mg related peaks in addition to organic-related peaks at 18, 19 and 29 Da. This would support the identification of peaks at 164 and 180 Da originating from spinochrome A. Contrastingly, the coincidence analysis for the peak at 224 Da reveals a much lower intensity peak at 29 Da, suggesting no COH^+^ ions are detected in coincidence events. This suggests the 224 Da peak is unlikely to be a fragment of spinochrome C, and may be unrelated to spinochrome A as the peak at 40 Da is at a similar intensity to the coincidence analysis of the other spinochrome A related ions. Overall, the coincidence analysis of each spinochrome A related peak reveals molecule fragments primarily through loss of CO, OH_3_ or CO_2_ molecules, which explains why these ions are frequently detected in biomineral APT datasets. This would also contrast the observation that C_x_H ions are primarily detected in APT analysis of organic molecules, and further highlights the differences between organic molecule incorporation into synthetic and biogenic calcite.

### 3.2 Spatially-resolved mass spectrometry

PHNQs are well-established molecules identified across several sea urchin species, but there are still questions regarding their spatial distribution from the micron-scale to the nanoscale within the biominerals. A further strength of APT is its capacity for mapping of the distribution of the identified ions in 3D at the nano-scale. Figure 10a provides the 3D distribution of a 350 x 50 x 50 nm^3^ volume in the purple region; of the ions at 208 Da (displayed as pink circles), at 7 Da (most likely Li^+^, orange circles), along with ions at 12 and 13 Da (Mg^2+^, blue dots) and 40 and 42 Da (most likely MgO^+^, green dots). The peak at 208 was determined as the best candidate for spinochrome A as it is unlikely to overlap with any known molecules, as is the case for peaks at 164 and 180 Da which could also be identified as monosaccharide moieties occluded within the purple region of the spines as discussed above. The 3D distributions of Mg^2+^ and MgO^+^ molecular ions are of particular interest as well due to their potential complexation with PHNQs^99^, with Li^+^ added to provide comparison for inorganic ions not reported to have association with PHNQs. Ca, C and O-containing ions relating to the calcite are omitted to focus on ions of interest.

**Figure 10.**
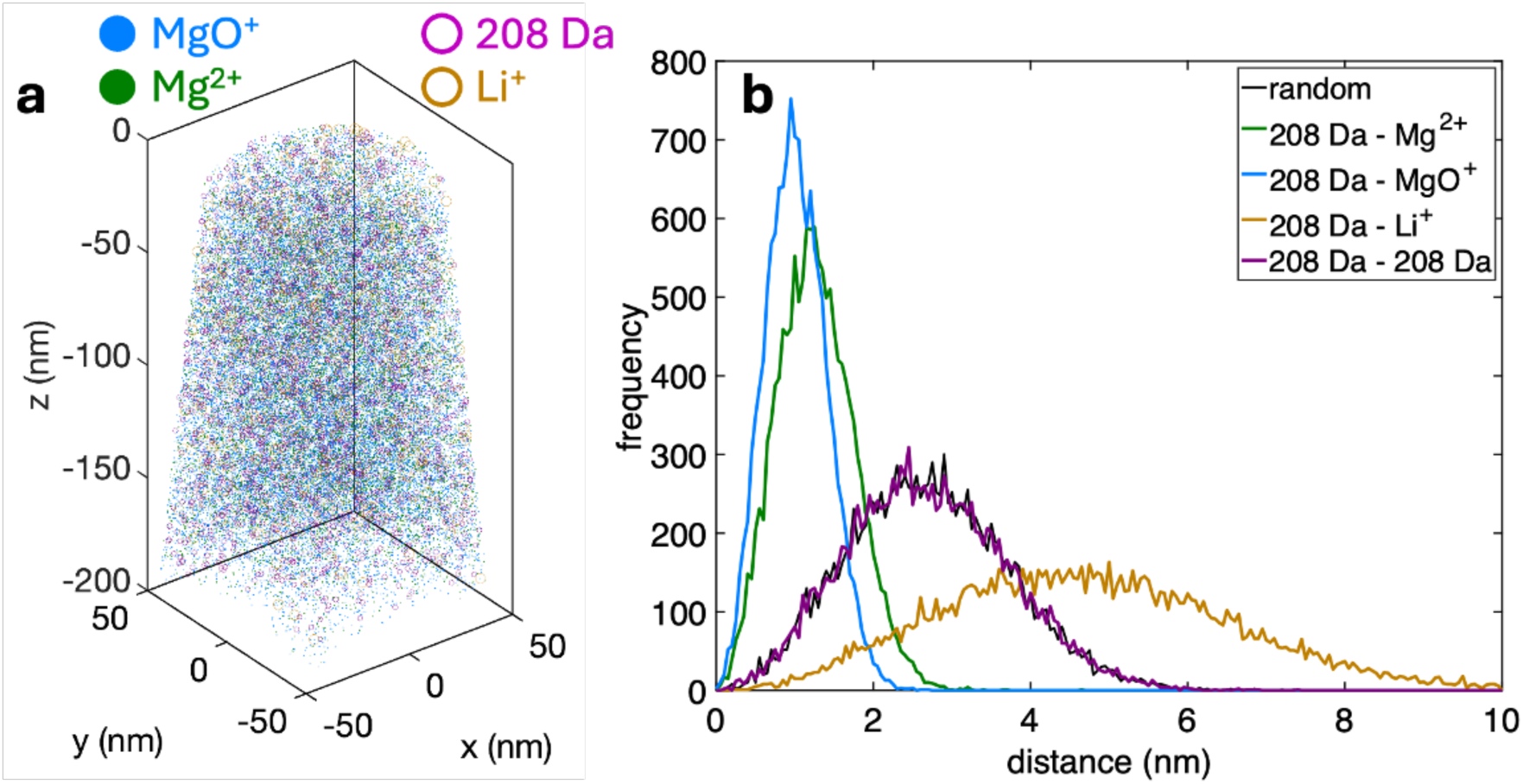
(a) 3D distribution in the purple region of the sea urchin spine of Mg^2+^, MgO^+^, Li^+^ and 208 Da (for visual clarity, no Ca-containing ions are shown, only 1 in 10 of the Mg-containing ions is displayed, and only 1 in 5 for the other ions). (b) First nearest neighbour distribution between the ions at 208 Da and the Mg^2+^, MgO^+^, Li^+^ and 208 Da ions along with the distribution obtained for a randomised dataset (black).

Based on Figure 10a, the distribution of all displayed ions appears homogeneous throughout the reconstructed data. To quantify if a species is homogenously distributed in APT, a common approach is to compare the histogram of the distance from one ion to its first nearest neighbour (1NN)^100,101^ in the experimental data and in a second dataset in which the mass-to-charge state ratios are assigned randomly but the spatial positions are kept the same. This approach is referred as random labelling and is commonly used to generate randomised APT datasets^101,102^ to assess clustering tendencies. Figure 10b plots these histograms for several ionic species and for the random distribution plotted as a black line that mimics a homogenous distribution. Here, a Kolmogorov–Smirnov test gives a p-value above 0.99 implying that the ions with a mass-to-charge of 208 Da that we associate to the spinochrome A are homogenously distributed throughout the analysed volume within the spatial resolution limits of APT^103^. The Li-Li and Mg-Mg are also both homogenously distributed (p value of 0.997 and 1 respectively, visualised in Figure S 7) throughout the analysed volume.

The distribution of the 1NN distance between an ion at 208 Da and respectively Mg^2+^/MgO^+^ ions (208 Da – Mg^2+^ and 208 Da – MgO^+^) or a Li^+^ ion (208 Da – Li^+^) are also plotted in Figure 10b. Here, the experimental distributions are not comparable to a random distribution with p-values below 10^-8^. The average shorter distance between 208 Da ions and a Mg^2+^/MgO^+^ ion in comparison to the randomised dataset (pink) implies a strong correlation of the distribution of Mg and fragments assumed to be from spinochrome A, and hence a co-segregation of these species. In contrast, the longer distance between 208 Da and Li^+^ reveals a strong anti-correlation between the 208 peak and the Li, suggesting these ions are spatially separated.

This strong association between the 208 Da fragment and Mg, in combination with the coincidence analysis which reported Mg and calcite-related peaks in events of the 208 Da ion (Figure 9b), indicate that spinochrome A is bound to Mg, consistent with previous reports for echinochrome A^99^. This is not the case for the other metallic impurities that were detected in the purple region of the spine, namely Si and Al (Figure S 8). Li is found in both regions, and known to affect the early development of sea urchins^104^, but its integration within the spine has not been studied so far. Here it is in addition spatially anti-correlated with spinochrome A and more concentrated in the non-pigmented tip of the spines (350 ppm in white and 250 ppm in purple). Additionally, in the white region (Figure S 9a-b, both Li and Mg ions are randomly distributed (p-values of 1 and 0.96, respectively –Figure S 9c). While it is possible that Mg concentration is influenced by a range of factors, including the location where specimens were prepared, the presence of spinochrome A pigments could influence Mg distribution and be responsible for higher concentrations recorded in the purple region.

## 4. Conclusion

Here, we use molecules responsible for the colouration of biomineral sea urchin spines to highlight the capabilities of APT to analyse organic materials and map their distribution at the nanoscale. New filtering techniques were introduced to determine the ions detected in coincidence with selected ions at specific mass-to-charge state ratios related to organic molecules, providing information on their fragmentation pathways under electric field conditions. Spinochrome molecules appear to fragment primarily through the loss of COH^+^ and OH_3_^+^ ions, however field conditions applied during APT analysis are not sufficiently high enough to cause complete fragmentation as is the case when these molecules are occluded within synthetic calcite, enabling their identification in the final mass spectrum of sea urchin spines. This may be due to an association with Mg in the calcite spines, as indicated by nearest neighbour analysis. This study provides a proof of concept that will help fast-track the necessary development of APT for spatially-resolved mass spectrometry of biological materials in 3D at the nanoscale. The methodology introduced can be applied to pre-existing APT datasets of biominerals, allowing for a re-analysis and potentially more accurate interpretation of the results. Novel generations of instruments may allow for improved signal to background ratios due to the implementation of the combination of voltage and laser pulsing for instance^105^, which may help detect lower intensity peaks that can then be used to perform more precise coincidence analysis. We expect that our work will also motivate the development of more advanced data processing techniques – possibly exploiting existing databases to help with fragment identification, along with machine-learning approaches commonly used in mass spectrometry, while also seeking theory and modelling work supported by quantum mechanical calculations.

