## Supplementary material for "Nanoscale identification of organic pigments in biogenic calcite by atom probe tomography"

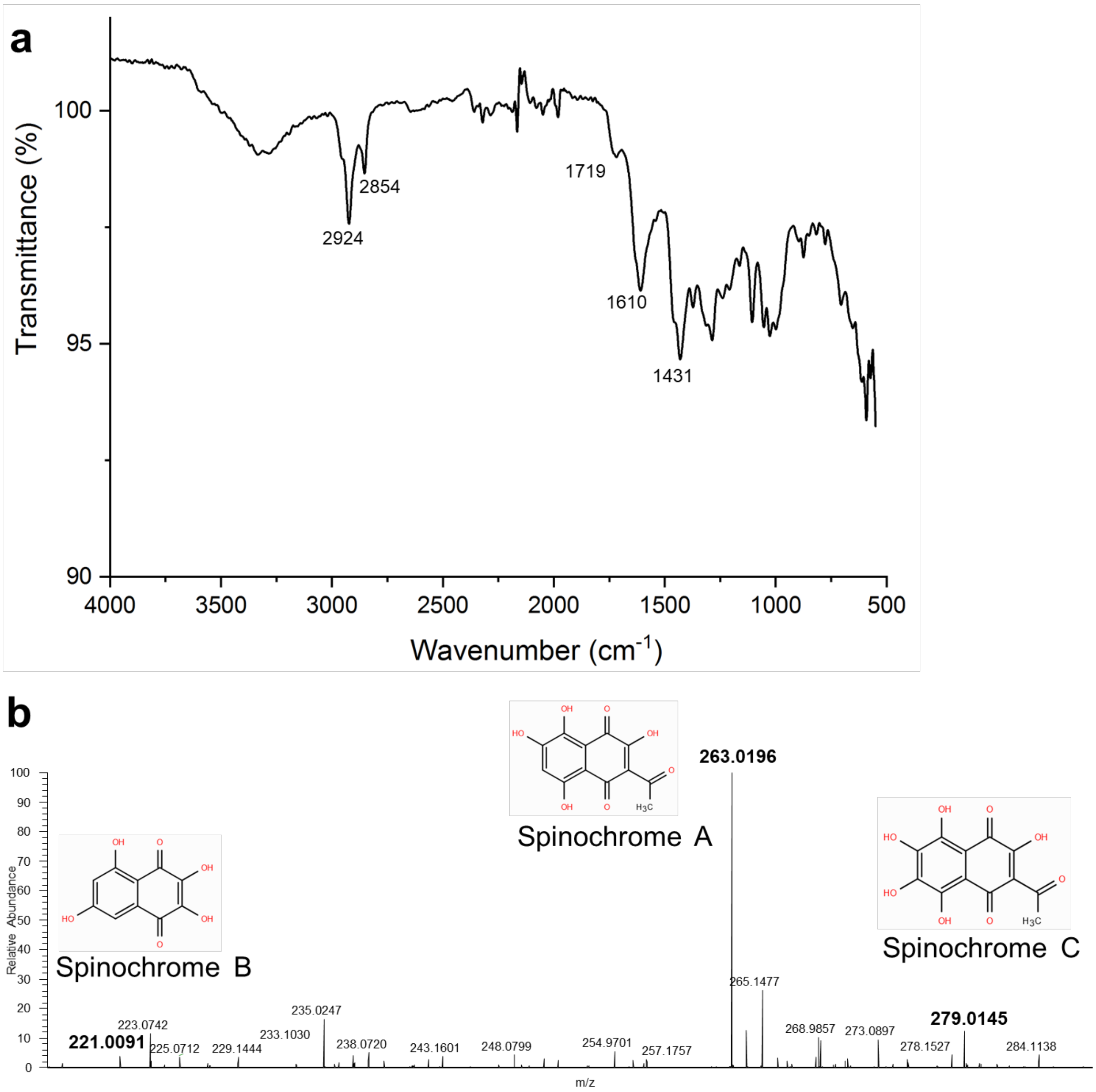


**Figure S 1**. a) FTIR spectra of the PHNQ extract from P. lividus displaying bands at 1719 and 1610 $cm^{-1}$ attributed to C=C and C=O stretching and C-O-H in plane of PHNQs[1]; the band at 1431 $cm^{-1}$ is attributed to protein backbone (N-H deformation in secondary amides)[2] and bands at 2924 and 2854 cm-1 to polysaccharides or/and lipopolysaccharides moieties[3], b) HR-MS spectra of the PHNQ extract from S. granularis showing the presence of mainly spinochrome A.

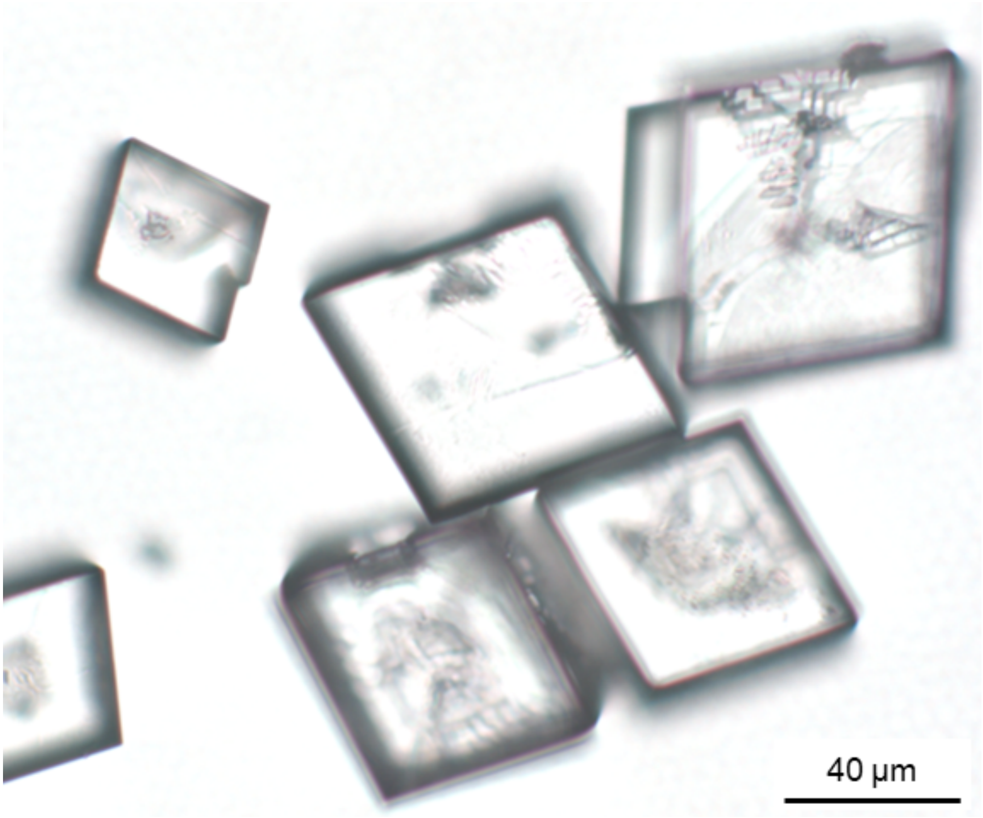


**Figure S 2.** Transmission optical microscope images of synthetic calcite crystals with rhombohedral morphology.

**Table S 1.** Peak identification for mass spectra from Figures 2a-b and 5a-b. Background represents origin of detected peak, from calcite (blue), PHNQ extract (purple), sea urchin spine (grey), and purple region of sea urchin spine only (red). Some peaks have multiple interpretations and could be overlaps from different ions with the same mass. Peaks that could not be identified are indicated by N/I (not identifiable).

| Mass-to-charge (Da) | Synthetic calcite | PHNQ doped calcite | Sea urchin spine (purple region) | Sea urchin spine (white region) |
|---|---|---|---|---|
| 1 | $H^+$ | $H^+$ | $H^+$ | $H^+$ |
| 2 | $H^{2+}$ | $H^{2+}$ | $H^{2+}$ | $H^{2+}$ |
| 3 | $H^{3+}$ | $H^{3+}$ | $H^{3+}$ | $H^{3+}$ |
| 7 | | | $Li^+$ | $Li^+$ |
| 12 | | $C^+$ | $Mg^{2+}/C^+$ | $Mg^{2+}/C^+$ |
| 12.5 | | | $Mg^{2+}$ | $Mg^{2+}$ |
| 13 | | | $Mg^{2+}/CH^+$ | $Mg^{2+}/CH^+$ |
| 13.5 | | | $Al^{3+}$ | $Al^{3+}$ |
| 14 | | | $Si^{2+}$ | $Si^{2+}$ |
| 15 | | $CH_3^+$/ $NH^+$ | $CH_3^+$/ $NH^+$ | $CH_3^+$/ $NH^+$ |
| 16 | $O^+$ | $O^+$ | $O^+$ | $O^+$ |
| 17 | $OH^+$ | $OH^+$ | $OH^+$ | $OH^+$ |
| 18 | $OH_2^+$ | $OH_2^+$ | $OH_2^+$ | $OH_2^+$ |
| 19 | $OH_3^+$ | $OH_3^+$ | $OH_3^+$ | $OH_3^+$ |
| 20 | $Ca^{2+}$ | $Ca^{2+}$ | $Ca^{2+}$ | $Ca^{2+}$ |
| 21 | $Ca^{2+}$ | $Ca^{2+}$ | $Ca^{2+}$ | $Ca^{2+}$ |
| 22 | $Ca^{2+}$ | $Ca^{2+}$ | $Ca^{2+}$ | $Ca^{2+}$ |
| 23 | $Na^+$ | $Na^+$ | $Na^+$ | $Na^+$ |
| 24 | $C_2^+$ | $C_2^+$ | $Mg^+/C_2^+$ | $Mg^+/C_2^+$ |
| 25 | | | $C_2H^+$ | $C_2H^+$ |
| 26 | | | $C_2H_2^+$ | $C_2H_2^+$ |
| 28 | $CO^+$ | $CO^+$ | $CO^+$ | $CO^+$ |
| 29 | $COH^+$ | $COH^+$ | $COH^+$ | $COH^+$ |
| 30 | $COH_2^+$ | $COH_2^+$ | $COH_2^+$ | $COH_2^+$ |
| 32 | $O_2^+$ | $O_2^+$ | $O_2^+$ | $O_2^+$ |
| 32.5 | | | $Mg_2O_2^+$ | $Mg_2O_2^+$ |
| 33 | | | $Mg_2O_2^+$ | $Mg_2O_2^+$ |
| 38 | | $O_2H_6^+$ | $C_2N^+/O_2H_6^+$ | $C_2N^+/O_2H_6^+$ |
| 39 | | | $C_2NH^+/K^+$ | $C_2NH^+/K^+$ |
| 40 | $Ca^+$ | $Ca^+$ | $MgO^+/C_2O^+/Ca^+$ | $MgO^+/C_2O^+/Ca^+$ |
| 40.5 | | | N/I | N/I |
| 41 | | | $MgO^+/C_2OH^+$ | $MgO^+/C_2OH^+$ |
| 42 | | | $MgO^+$ | $MgO^+$ |
| 43 | | | $CONH^+$ | $CONH^+$ |
| 44 | $CO_2^+$ | $CO_2^+$ | $CO_2^+$ | $CO_2^+$ |
| 45 | $CO_2H^+$ | $CO_2H^+$ | $CO_2H^+$ | $CO_2H^+$ |
| 46 | | | $CO_2H_2^+$ | $CO_2H_2^+$ |
| 47 | | | $PO^+$ | $PO^+$ |
| 48 | $Ca_2O^+$ | $Ca_2O^+$ | $Ca_2O^+$ | $Ca_2O^+$ |
| 49 | $Ca_2O^+$ | $Ca_2O^+$ | $Ca_2O^+$ | $Ca_2O^+$ |

| | | | | |
|---|---|---|---|---|
| 50 | $Ca_2O^+$ | $Ca_2O^+$ | $Ca_2O^+$ | $Ca_2O^+$ |
| 54 | | | $C_2ON^+$ | $C_2ON^+$ |
| 56 | $CaO^+$ | $CaO^+$ | $CaO^+$ | $CaO^+$ |
| 57 | $CaO^+$ | $CaO^+/C_3OH_5^+$ | $CaO^+/C_3OH_5^+$ | $CaO^+/C_3OH_5^+$ |
| 58 | $CaO^+$ | $CaO^+$ | $CaO^+$ | $CaO^+$ |
| 59 | $CaO^+$ | $CaO^+/C_2ONH_5^+$ | $CaO^+/C_2ONH_5^+$ | $CaO^+/C_2ONH_5^+$ |
| 60 | $CO_3^+$ | $CO_3^+$ | $CO_3^+$ | $CO_3^+$ |
| 61 | | | $C_5H^+/C_2O_2H_3^+$ | $C_5H^+/C_2O_2H_3^+$ |
| 62 | | | N/I | N/I |
| 62.5 | | | N/I | N/I |
| 63 | | | N/I | N/I |
| 64 | | | $C_5H_4^+$ | $C_5H_4^+$ |
| 65 | | | $C_5H_5^+$ | $C_5H_5^+$ |
| 68 | | | $C_3O_2^+$ | $C_3O_2^+$ |
| 69 | $Ga^+$ | $Ga^+$ | $Ga^+$ | $Ga^+$ |
| 70 | $Ca_2CO_3^{2+}$ | $Ca_2CO_3^{2+}$ | $Ca_2CO_3^{2+}$ | $Ca_2CO_3^{2+}$ |
| 71 | $Ca_2CO_3^{2+}$ | $Ca_2CO_3^{2+}$ | $Ca_2CO_3^{2+}$ | $Ca_2CO_3^{2+}$ |
| 72 | $CaO_2^+$ | $CaO_2^+$ | $CaO_2^+$ | $CaO_2^+$ |
| 73 | | | N/I | N/I |
| 74 | $Ca_4CO^{2+}$ | $Ca_4CO^{2+}$ | $Ca_4CO^{2+}$ | $Ca_4CO^{2+}$ |
| 75 | | $C_2O_2NH_5^+$ | $C_2O_2NH_5^+$ | $C_2O_2NH_5^+$ |
| 76 | $CO_4^+$ | $CO_4^+$ | $CO_4^+$ | $CO_4^+$ |
| 77 | | | $C_6H_5^+$ | $C_6H_5^+$ |
| 78 | | | $C_6H_6^+$ | $C_6H_6^+$ |
| 79 | | $C_6H_7^+$ | $C_6H_7^+$ | $C_6H_7^+$ |
| 80 | | | $PO_3H^+$ | $PO_3H^+$ |
| 81 | | | $PO_3H_2^+$ | $PO_3H_2^+$ |
| 82 | | $C_4O_2H_2^+$ | $C_4O_2H_2^+$ | $C_4O_2H_2^+$ |
| 83 | | | N/I | N/I |
| 84 | | | $MgCO_3^+$ | $MgCO_3^+$ |
| 85 | | | $C_7H^+$ | $C_7H^+$ |
| 88 | | | $Sr^+/C_6O^+$ | $Sr^+/C_6O^+$ |
| 89 | | | $C_6OH^+$ | $C_6OH^+$ |
| 90 | | | $C_6OH_2^+$ | $C_6OH_2^+$ |
| 92 | | | N/I | N/I |
| 93 | | | N/I | N/I |
| 96 | | | N/I | N/I |
| 97 | | | $C_8H^+$ | $C_8H^+$ |
| 98 | | | $C_8H_2^+$ | $C_8H_2^+$ |
| 99 | | | $C_8H_3^+$ | $C_8H_3^+$ |
| 100 | $CaCO_3^+$ | $CaCO_3^+$ | $CaCO_3^+$ | $CaCO_3^+$ |
| 101 | $CaCO_3^+$ | $CaCO_3^+$ | $CaCO_3^+$ | $CaCO_3^+$ |
| 102 | $CaCO_3^+$ | $CaCO_3^+$ | $CaCO_3^+$ | $CaCO_3^+$ |
| 103 | | | $C_8H_7^+$ | $C_8H_7^+$ |
| 104 | $C_2O_5^+$ | $C_2O_5^+$ | $C_2O_5^+$ | $C_2O_5^+$ |
| 112 | $Ca_2O_2^+$ | $Ca_2O_2^+$ | $Ca_2O_2^+$ | $Ca_2O_2^+$ |
| 113 | | | N/I | N/I |
| 118 | | | N/I | N/I |
| 119 | | | N/I | N/I |
| 120 | | | N/I | N/I |

| | | | | |
|---|---|---|---|---|
| 128 | | | $MgC_2O_4^+$ | $MgC_2O_4^+$ |
| 129 | | | $MgC_2O_4^+$ | $MgC_2O_4^+$ |
| 130 | | | $MgC_2O_4^+$ | $MgC_2O_4^+$ |
| 136 | | | $C_5O_4H_{12}^+$ | $C_5O_4H_{12}^+$ |
| 137 | | | $C_5O_4H_{13}^+$ | |
| 138 | | | $C_5O_4H_{14}^+$ | |
| 140 | | | $MgC_3O_4^+$ | $MgC_3O_4^+$ |
| 144 | $CaC_2O_4^+$ | $CaC_2O_4^+$ | $CaC_2O_4^+$ | $CaC_2O_4^+$ |
| 164 | | | $C_9O_3H_8^+$ | |
| 172 | | | $MgC_3O_6^+$ | $MgC_3O_6^+$ |
| 173 | | | $MgC_3O_6^+$ | $MgC_3O_6^+$ |
| 174 | | | $MgC_3O_6^+$ | $MgC_3O_6^+$ |
| 180 | | | $C_9O_4H_8^+$ | |
| 188 | $CaC_3O_6^+$ | $CaC_3O_6^+$ | $CaC_3O_6^+$ | $CaC_3O_6^+$ |
| 208 | | | $C_{10}O_5H_8^+$ | |
| 216 | | | $MgC_4O_8^+$ | $MgC_4O_8^+$ |
| 224 | | | $C_{10}O_6H_8^+$ | |
| 232 | | $C_{12}O_5H_8^+$ | | |
| 254 | | | $C_{11}O_6H_{10}^+$ | |

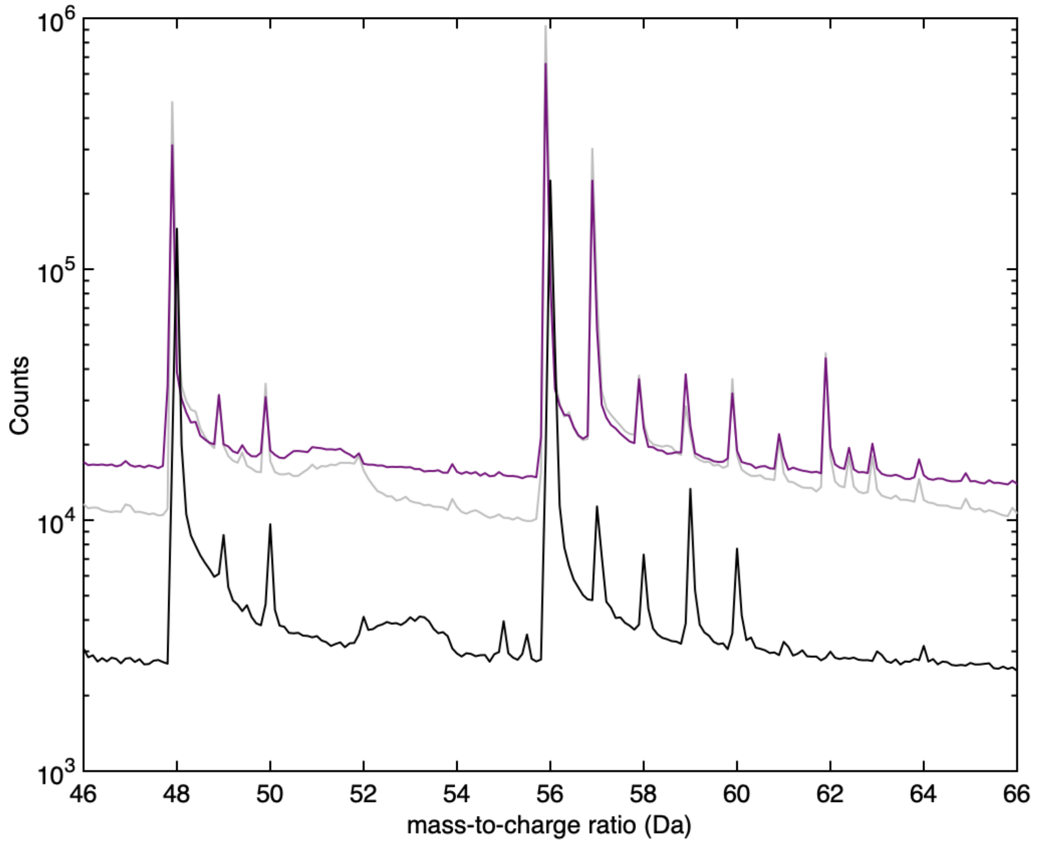


**Figure S 3.** Close up in the range 46–66 Da of the mass spectra of the pure calcite (black) and sea urchin spine in the white region (grey) and purple region (purple).

**Table S 2.** List of isotopes and relative abundance for $Ca^+$ and $MgO^+$.

| | $Ca^+$ | | $MgO^+$ | |
|---|---|---|---|---|
| Mass-to-charge (Da) | natural abundance | relative amplitude | natural abundance | relative amplitude |
| 40 | 0.969 | 1 | 0.789 | 1 |
| 41 | | | 0.100 | 0.127 |
| 42 | 0.007 | 0.007 | 0.112 | 0.139 |
| 43 | 0.001 | 0.001 | | 0.002 |
| 44 | 0.020 | 0.021 | | |
| 48 | 0.028 | 0.029 | | |

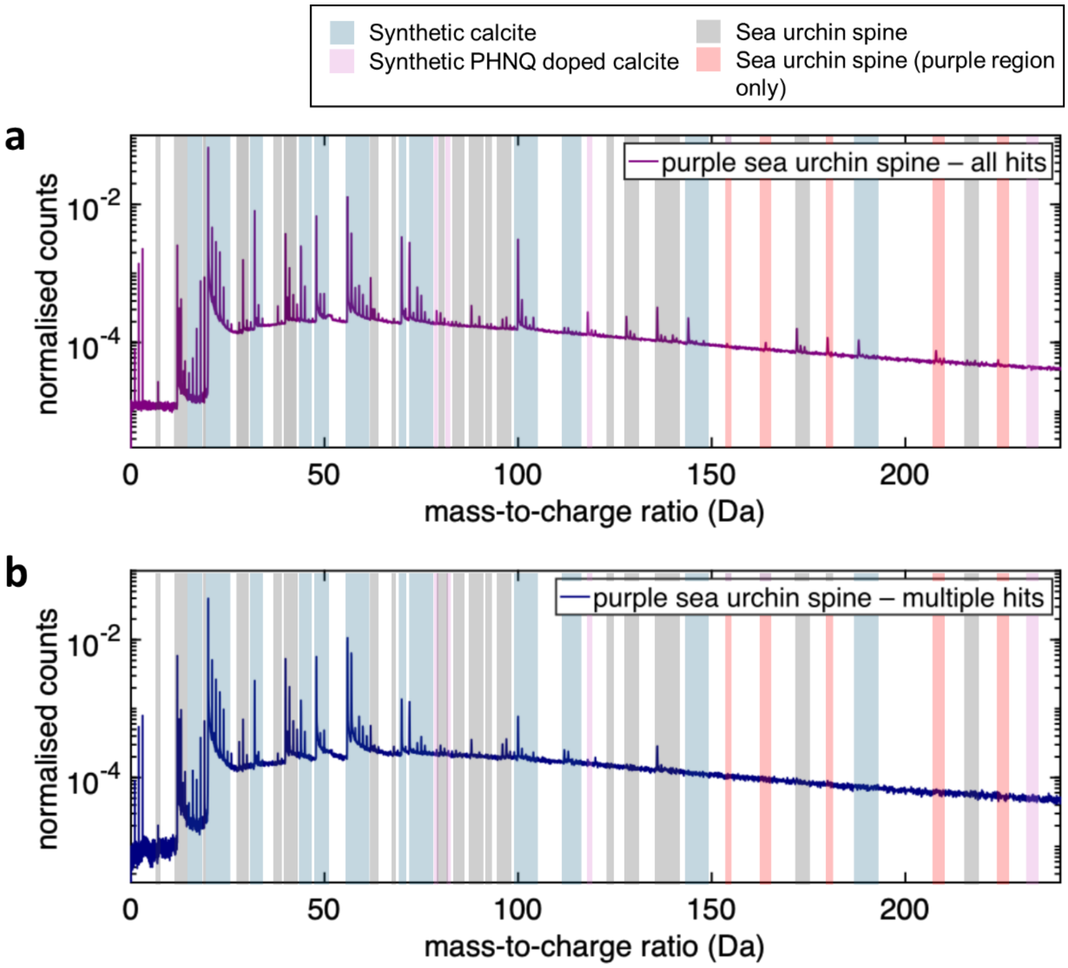


**Figure S 4.** Mass spectrum for the purple region of the sea urchin spine (a) for all hits and (b) just for ions detected as part of multiple hits.

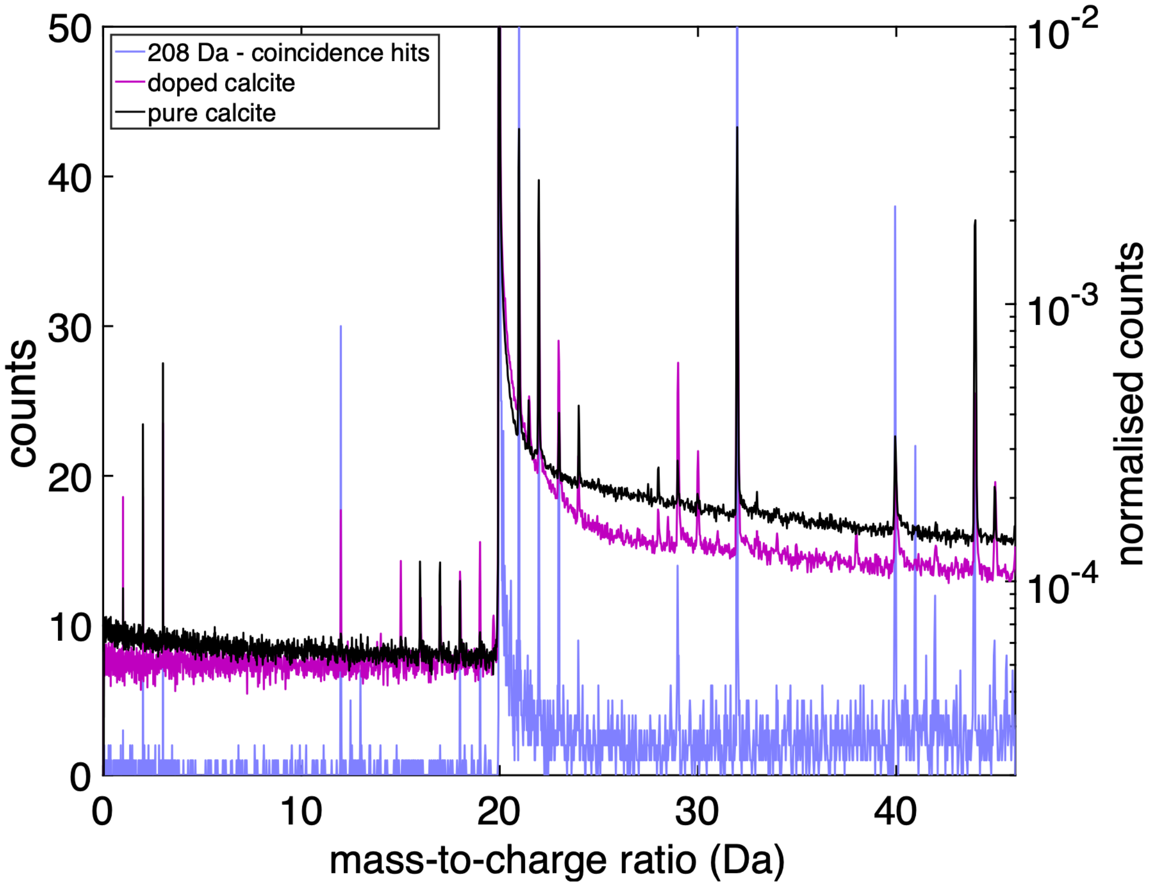


**Figure S 5.** Mass spectra for the coincidence analysis for the range at 208 Da of the purple region of the sea urchin spine in lilac, plotted in linear scale on the left y-axis; and for the pure and doped calcite plotted in log scale on the right y-axis, normalised to the total number of ions in the dataset.

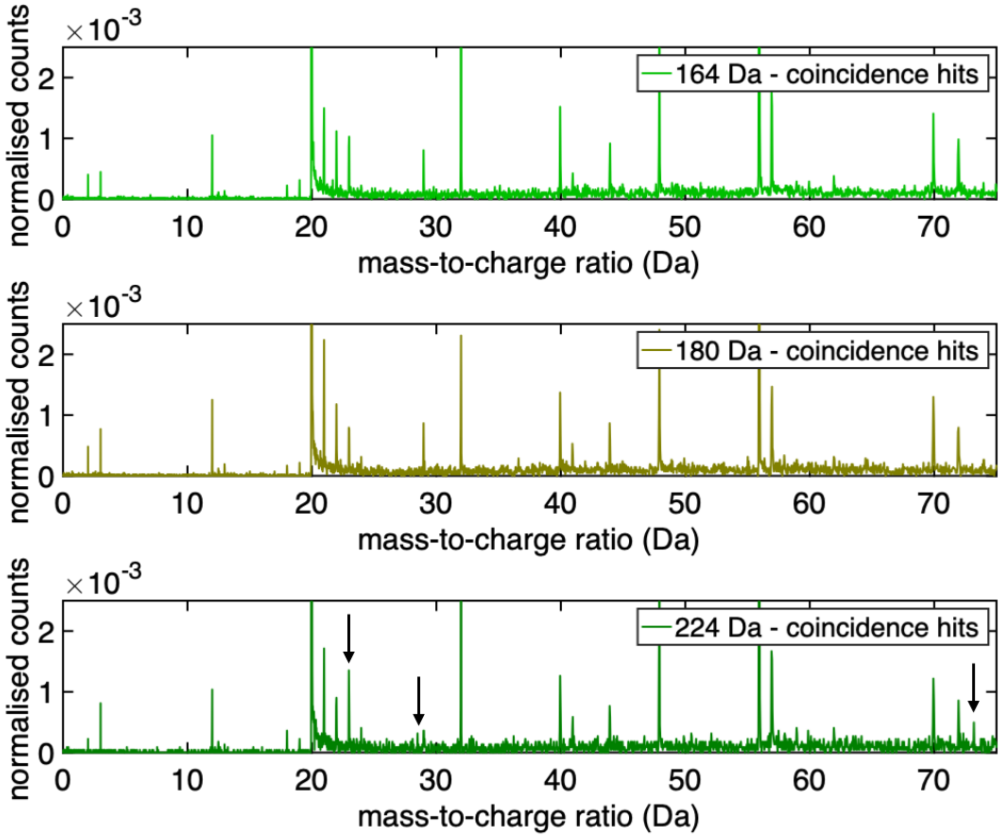


**Figure S 6.** Mass spectrum of coincidence analysis for ions at mass ranges 164, 180 and 224 Da. Differences in the coincidence analysis of 224 Da to other ions is indicated by black arrows.

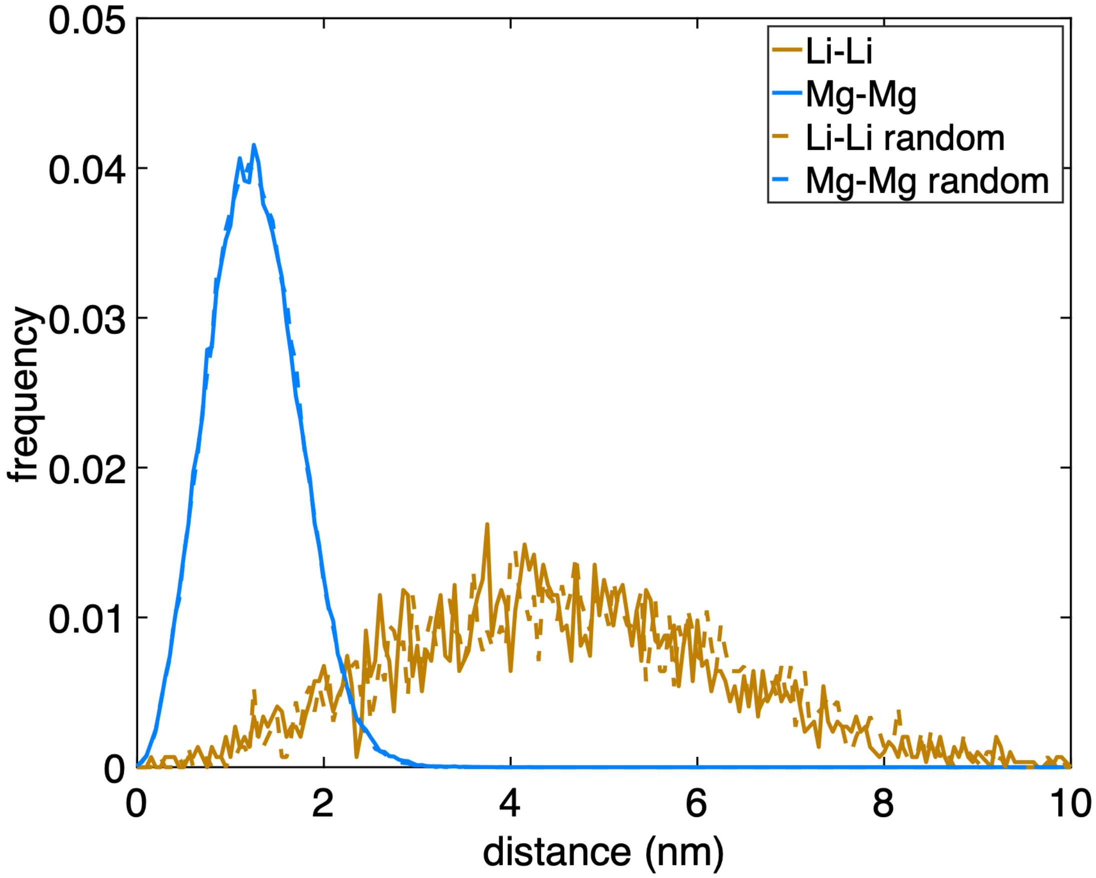


**Figure S 7.** First nearest neighbour distribution between two ions in the purple spine region at 7 Da (Li) in blue and two ions at 12 Da (Mg) in blue. Dashed lines indicate randomised dataset.

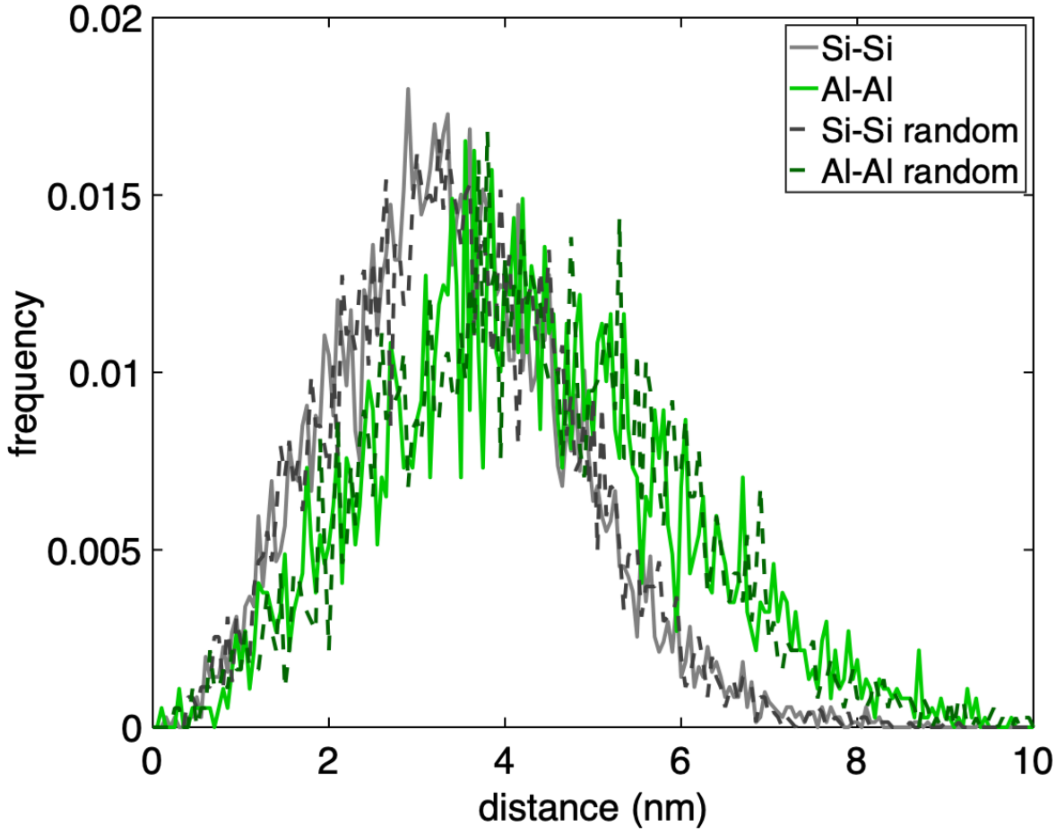


**Figure S 8.** First nearest neighbour distribution between two ions at 14 Da (Si) in grey and two ions at 13.5 Da (Al) in green. Dashed lines indicate randomised dataset.

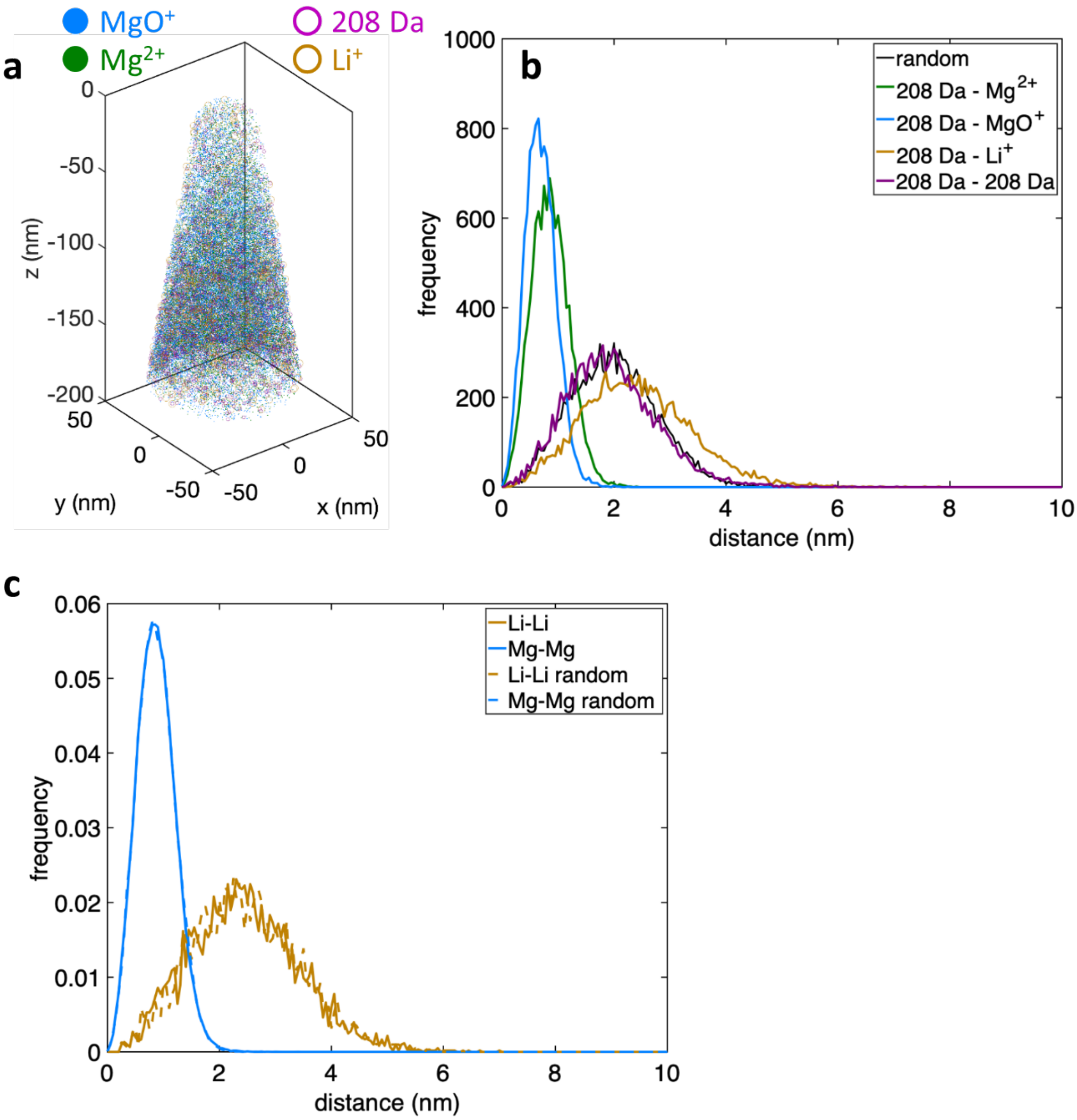


**Figure S 9.** (a) 3D distribution in the white region of the sea urchin spine of $Mg^{2+}$, $MgO^{+}$, $Li^{+}$ and 208 Da (for visual clarity, no Ca-containing ions are shown, only 1 in 10 of the Mg-containing ions is displayed, and only 1 in 5 for the other ions). (b) First nearest neighbour distribution between the ions at 208 Da and the $Mg^{2+}$, $MgO^{+}$, $Li^{+}$ and 208 Da ions along with the distribution obtained for a randomised dataset (black). (c) First nearest neighbour distribution between two ions at 7 Da (Li) in blue and two ions at 12 Da (Mg) in blue. Dashed lines indicate randomised dataset.
